# Affinity-tag-based microfluidic protein isolation enables high-resolution Cryo-EM from minimal starting material

**DOI:** 10.64898/2026.05.20.726462

**Authors:** M. Zimmermann, D. E. Schneider, L. Rima, T. Clairefeuille, R. Thoma, M. E. Lauer, T. Braun

## Abstract

Cryo-EM is central to high-resolution structure determination, but conventional sample preparation consumes substantial amounts of protein and can destabilize sensitive complexes. Microfluidic approaches can miniaturize and accelerate these workflows while retaining the particle numbers required for single-particle analysis. Here, we present a generalized microfluidic isolation strategy that captures proteins directly from cell lysates or *in vitro* translation reactions using genetically encoded tags. The platform supports both affinity-based (ALFA-nanobody) and covalent (SpyTag3/SpyCatcher3) capture. Specificity derives from two sequential, orthogonal steps: (i) tag-mediated capture and concentration of the target protein on beads, and (ii) photoelution from the bead surface. By confining the sample to nanoliter volumes and minimizing transport, the approach suppresses non-specific carryover. Importantly, the isolation workflow is coupled to grid preparation, enabling preparation of three cryo-EM or negative-stain grids from a 25 nL eluate. Using this workflow, we isolated *E. coli* ferritin A, β-galactosidase, and *P. aeruginosa* VgrG1 from cell lysate, obtaining reconstructions at 1.9–2.6 Å resolution with B-factors comparable to optimized conventional workflows. We further isolated ferritin A from an *in vitro* translation reaction, yielding a 2.04 Å reconstruction. A standardized 50 µL starting volume enabled more than 16 independent microfluidic isolations. Tag-based microfluidic isolation thus provides a broadly applicable route to cryo-EM sample preparation, reducing sample volumes more than 1000-fold, reducing preparation times more than 3-fold for lysate-based workflows and up to 10-fold when combined with cell-free expression, and enabling high-throughput structural screening directly from *in vitro* translation reactions.

**Significance Statement:** Cryo-electron microscopy (cryo-EM) enables protein structure determination at atomic resolution, but conventional sample preparation requires large amounts of purified protein and laborious workflows that can damage fragile complexes. We present a microfluidic platform that captures tagged proteins directly from minute amounts of crude lysates or cell-free synthesis reactions and deposits them onto cryo-EM grids in a single integrated workflow. Using this approach, we determine three protein structures at high resolution. By reducing sample volumes by roughly three orders of magnitude, the platform expands cryo-EM to proteins inaccessible to conventional workflows and provides a foundation for automated, high-throughput structural analysis.

## 1 Introduction

Elucidating high-resolution protein structures is essential for understanding molecular mechanisms, enzymatic functions and for enabling structure-based drug design. Over the past decade, cryogenic electron microscopy of vitrified specimens (cryo-EM) has undergone a rapid technological maturation (1, 2), and single-particle analysis (SPA) has become a method of choice for high-resolution structure determination of protein complexes. This progress has been driven by advances on multiple fronts: improved hardware, in particular the introduction of direct electron detectors, enabling higher-quality data acquisition (3); a high degree of automation in data collection (4); and substantial developments in image processing and reconstruction algorithms (5, 6). In parallel, atomic model building has been accelerated by machine-learning-based tools that predict protein structures from amino acid sequences (e.g. AlphaFold (7) for initial modeling) and by methods that assist model building directly from reconstructed cryo-EM maps such as ModelAngelo (8). As a result, near-atomic resolution structures (<3 Å) of many protein complexes can now be obtained routinely.

Despite these advances, sample preparation remains a major bottleneck in the cryo-EM workflow (9–12). The overall preparation pipeline typically comprises three major steps: (i) production of sufficient amounts of the target protein, for example by recombinant overexpression; (ii) purification of the target protein; and (iii) cryo-EM grid preparation. All of these steps present distinct technical challenges and require substantial amounts of time, material, and expertise. Notably, only 10^4^–10^6^ particles are typically required for high-resolution structure determination by single-particle cryo-EM, with flexible or heterogeneous assemblies requiring more particles to achieve high-resolution reconstructions. Handling such small amounts of material makes microfluidic approaches particularly attractive for cryo-EM sample preparation.

For classical protein purification workflows, liters of cell culture are required to obtain a sufficient amount of purified protein, as typical protein concentrations used for single-particle cryo-EM are in the low micromolar range. This is often difficult to achieve for low-abundance proteins, unstable complexes, or proteins that cannot be overexpressed in cellular systems due to toxic effects. In addition, state-of-the-art purification procedures remain time-consuming and typically rely on one or more chromatographic steps. Such chromatographic workflows can be detrimental to fragile protein complexes due to dilution, shear forces, repeated surface interactions and prolonged processing times. In contrast, batch purification approaches can be gentler and allow pre-concentration of target proteins, but they often suffer from insufficient purity due to the co-release of non-specifically bound contaminant proteins by competitive elution.

Furthermore, conventional grid preparation requires microliter volumes of sample per grid and harsh blotting steps can damage proteins through shear forces and repeated exposure to the air-water interface (11, 13–15). In addition, the vast majority of the applied sample is lost during blotting (12, 16), dramatically increasing the effective protein consumption per cryo-EM grid. Although classical blotting-based preparation is a well-established, simple and widely accessible method, it remains a largely trial-and-error based process that further exacerbates material consumption and limits throughput.

To address these limitations, we developed a semi-automated microfluidic protein isolation and cryo-EM preparation platform that combines tag-based batch capture and photoelution to ensure highly specific release of the target protein while minimizing co-elution of non-specifically bound contaminants. Subsequently, the direct, blotting-free deposition onto cryo-EM grids enables vitrification and high-resolution structural analysis from minimal starting material. In contrast to conventional chromatographic workflows, our approach maintains the target protein in confined nanoliter volumes throughout purification and grid preparation, which helps preserve weak protein-protein interactions. Furthermore, our method reduces the initial sample volume by more than three orders of magnitude and shortens purification and grid preparation times by more than fivefold.

Building on our previous work (17, 18), we extend the platform to a generalized, affinity-tag-based isolation strategy, removing the need for protein-specific capture reagents and improving robustness across different targets. Using ALFA-nanobody- and SpyCatcher3-based capture (19, 20), we demonstrate microfluidic isolation and direct cryo-EM preparation from as little as 10 µL of cell lysate or 50 µL of an *in vitro* translation reaction. These workflows yield high-resolution reconstructions with Rosenthal-Henderson B-factors averaging 54.6 Å^2^ for proteins isolated from cell lysates and 67.4 Å^2^ for proteins isolated directly from *in vitro* translation reactions. These values are within or below the range typically reported for well-optimized conventional grid preparation workflows.

Beyond enabling protein isolation from minimal starting material, the microfluidic workflow also provides an experimental framework to examine how capture chemistry, tag design, and microfluidic sample handling influence cryo-EM sample preparation and structural data quality.

## 2 Results

### 2.1 Generalized non-chromatographic microfluidic isolation of tagged proteins

We established a generalized, non-chromatographic workflow for microfluidic batch isolation of tagged proteins, enabling direct cryo-EM sample preparation from complex mixtures. To assess the generality and performance of the approach, we evaluated four tagging strategies: three non-covalent affinity tags (His-tag (31), FLAG-tag (32), and ALFA-tag (19)) and one covalent capture system (SpyTag3/SpyCatcher3 (20)). In addition, we tested the compatibility of the workflow with GFP-tagged proteins (33, 34), a widely used fluorescent reporter, as an exploratory proof-of-concept. These five tagging strategies and their corresponding capture reagents are summarized in Table 1. All capture strategies were implemented on superparamagnetic beads within a microcapillary system, and elution was achieved by photocleavage of a covalent linker that connects the capture reagent to the bead surface.

**Table 1.** Benchmarking of affinity tags for microfluidic protein isolation. Summary of tagging strategies tested in this study, including tag size, capture reagents, bead functionalization strategy, and experimental outcome. Details of the immobilization chemistry are provided in Supplementary Information SI-1A.

| Tag | Size [aa] | Capture reagent | Immobilization strategy | Outcome |
| --- | --- | --- | --- | --- |
| His<sub>6</sub> | 6 | Ni-NTA crosslinkers | Biotinylation via primary amine group on the NTA crosslinker | Good yield, but released NTA species and unspecific binding interfered with cryo-EM analysis |
|  | 6 | Aptamer | Biotinylation via primary amine group on the aptamer | Insufficient affinity of tested aptamers (see SI-4) |
|  | 6 | Anti-His antibody / Fab | Biotinylation via primary amines on antibody / Fab | Insufficient yield |
| FLAG | 8 | Anti-FLAG antibody / Fab | Biotinylation via primary amines on antibody / Fab | Functionalization reduced binding efficiency |
| ALFA | 15 | ALFA nanobody | DBCO functionalization via engineered cysteine followed by biotinylation with photocleavable linker | Reproducible and robust isolation from cell lysates and IVT |
| SpyTag3 | 16 | SpyCatcher3 | DBCO functionalization via engineered cysteine followed by biotinylation with photocleavable linker | Reproducible and robust isolation from cell lysates |
| GFP | 238 | Anti-GFP nanobody | Biotinylation via primary amines on nanobody | Successful capture; structural instability of tested constructs |

The generalized microfluidic batch isolation workflow is depicted in Figure 1 and builds on our previous work (17, 18). In brief, superparamagnetic beads, functionalized with capture reagents via a photocleavable linker, are immobilized inside a fused silica capillary using a magnetic trap that generates strong magnetic field gradients. Target proteins are captured from cell lysates or *in vitro* translation reactions either by flowing the sample past the immobilized beads or by pre-incubating the beads with the sample in a reaction vessel prior to uptake into the capillary. Following washing steps to remove unbound material, the beads are confined between two air bubbles, forming a volume of ~30 nL. Upon UV-illumination, the photocleavable linker is cleaved, releasing the target protein together with the bound capture reagent. This eluate is then deposited directly onto cryo-EM grids in a blotting-free manner for vitrification. Importantly, the excess liquid is reaspirated into the capillary prior to vitrification, which allows the preparation of multiple grids from one isolation. More details of the microfluidic setup and workflow are provided in Supplementary Information SI-2.

**Figure 1:**
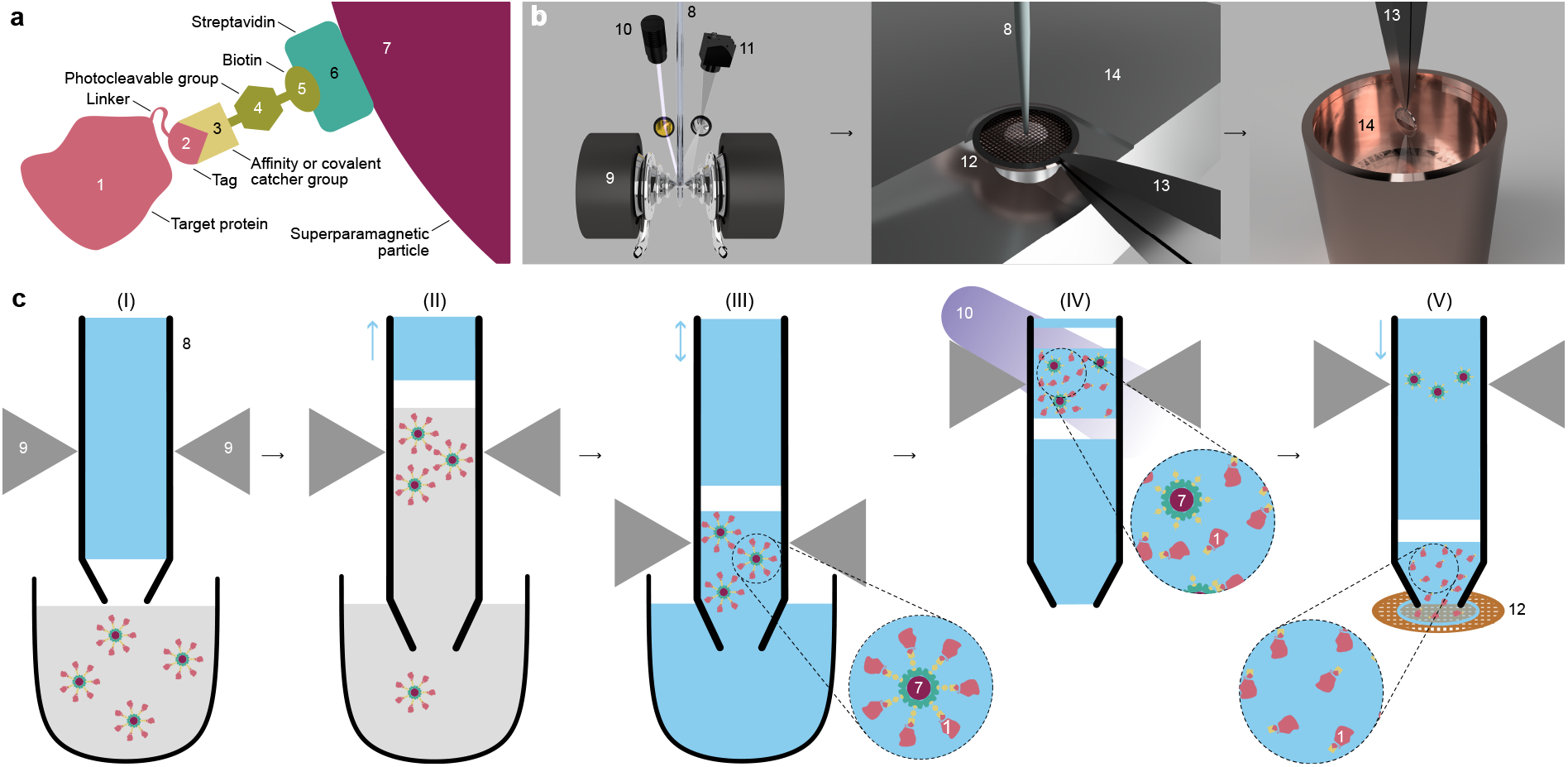
Microfluidic protein isolation and cryo-EM sample preparation workflow. The target protein is isolated through two sequential, specific steps: first, tag-based immobilization; second, specific photocleavage of the capture molecule. Notably, the same nozzle is used for both protein isolation and grid preparation. The sample is displaced only minimally within the microfluidic system, while the robotics enable relative movement of the nozzle and the magnetic trap. **a)** Composite system used for protein capture. The target protein (1), carrying a tag (2), is captured by an affinity or covalent binder (3). This binder is linked via a crosslinker containing a photocleavable moiety (4) and a biotin group (5), which binds strongly to streptavidin-functionalized (6) superparamagnetic beads (7). **b)** Hardware for protein isolation (left to right). For protein purification, the beads are trapped within a fused-silica microcapillary (8) using a water-cooled electromagnetic trap (9). A UV-source (10) enables photocleavage, and the process is monitored via a camera (11). The EM grid (12) is held by tweezers (13) on a temperature-controlled stage (14) during sample deposition. After writing a thin liquid layer, the grid is horizontally withdrawn, rotated by 90°, and plunged into a liquid ethane reservoir. **c)** Schematic of the microfluidic workflow for protein extraction and cryo-EM sample preparation: *(I)* insertion of the capillary into the magnetic trap and aspiration of a small air bubble; *(II)* uptake and trapping of functionalized beads, concentrating the target protein; *(III)* washing and positioning of the beads at the capillary tip, confined by an air bubble to a minimal buffer volume of ~30 nL; *(IV)* UV-induced photocleavage to elute bound proteins while the beads remain trapped; *(V)* deposition (“writing”) of the eluted protein onto a cryo-EM grid. Blue arrows alongside the capillary indicate pump direction.

Using this workflow, up to three cryo-EM grids can be prepared from a single isolation cycle, or alternatively two cryo-EM grids and one negative-stain grid. Negative-stain EM provides a convenient and fast assessment of sample quality and particle integrity after isolation, bypassing the need for conventional concentration measurements or gel electrophoresis, which are not feasible at nanoliter elution volumes.

Photoelution from the bead surface releases the captured protein-binder complex specifically, in contrast to competitive elution, which can liberate a broader spectrum of non-specifically adsorbed contaminants, which is particularly relevant under the high surface-to-volume ratios of microfluidic systems.

Robust capture reagents and appropriate bead functionalization are critical for the performance of the workflow, and tagging strategies that perform well in chromatographic workflows are not necessarily optimal for confined microfluidic batch isolation (see subsection 2.2). Details regarding the crosslinking chemistry and functionalization strategies are provided in Supplementary Information SI-1A.

### 2.2 Tag performance in batch capture differs from chromatographic workflows

His-tags are among the most widely used affinity tags in chromatographic purification workflows, often combining immobilized metal ion affinity chromatography (IMAC) and size exclusion chromatography (SEC). To evaluate their suitability for microfluidic batch isolation, we tested several strategies for capturing His-tagged proteins (see Table 1). These strategies can be broadly divided into two categories.

First, we employed photocleavable NTA-derivatives that allow immobilization of His-tagged proteins through metal chelation, analogous to classical IMAC. NTA-functionalized beads showed substantial non-specific binding even in the presence of imidazole or Tween, and target yields were typically too low for cryo-EM. Trivalent NTA improved binding strength but did not resolve the specificity problem, and photocleaved NTA derivatives interfered with downstream imaging (SI-3A).

Second, we explored capture strategies using other His-tag-binding capture reagents, including antibodies that specifically recognize His-tag epitopes and their Fab fragments. While these reagents showed reasonable equilibrium binding affinities (*K*_*D*_ ~ 100 nmol dm^−3^), the resulting capture efficiency in the microfluidic batch workflow remained limited. We attribute this to rapid binding kinetics with fast dissociation rates, which can lead to partial loss of target proteins during washing steps of our workflow (see SI-3A and SI-4 for additional details).

We also evaluated FLAG-tag-based capture for microfluidic protein isolation. Under the tested conditions, FLAG-tag capture resulted in comparatively low yields that were insufficient for routine cryo-EM data collection. A likely culprit for the limited performance of FLAG-based capture is the amine-targeting biotinylation strategy. Random lysine biotinylation can occur close to the antigen-binding site and thereby reduce binding efficiency. This is a particular limitation for antibody-based reagents, where site-directed functionalization is rarely possible.

Our observations indicate that capture chemistry well-suited to chromatography does not transfer directly to microfluidic batch workflows; further optimization would be required. In this study, we did not pursue further optimization of the His- and FLAG-based workflows and instead focused on capture strategies with well-defined and accessible functionalization sites, such as ALFA-nanobody, SpyTag/SpyCatcher3, and GFP-nanobody systems.

### 2.3 Isolation of Ferritin A using ALFA-tag and SpyTag3

To demonstrate robust microfluidic isolation and cryo-EM preparation, we analyzed two Apoferritin constructs: FtnA carrying an N-terminal His_6_-SpyTag3 fusion (Spy-ApoF) and FtnA carrying an N-terminal His_6_-ALFA tag (ALFA-ApoF). Both proteins were expressed in *E. coli*, and clarified cell lysates were used directly for microfluidic isolation. All protein constructs are described in more detail in Supplementary Information SI-1B.

Apo-ferritin is one of the most widely used benchmark proteins for cryo-EM grid preparation and data processing. At the same time, its 24-mer assembly represents a demanding test case for our workflow, as efficient photoelution must release a large multimeric complex from the capture reagents without disrupting the particle. Therefore, this provides a stringent test of the efficiency of the photoelution step.

Protein expression and accessibility of the affinity tags were verified using complementary biochemical assays, including fluorescence labeling of SpyTag3 constructs, pull-down experiments, and bio-layer interferometry measurements (see Supplementary Information SI-4).

Microfluidic isolation of Spy-ApoF and ALFA-ApoF was subsequently performed directly from clarified lysates. Efficient photoelution produced sufficient protein for cryo-EM grid preparation, demonstrating that the workflow can reliably isolate multimeric protein assemblies. Initial experiments revealed aggregation of Apo-ferritin particles on cryo-EM grids, which we traced to residual metal-ion interactions mediated by the His_6_-tag. Addition of 1 mmol dm^−3^ EDTA to the buffer eliminated this effect and resulted in well-dispersed particles.

Cryo-EM data was collected on a FEI Titan Krios G4 and processed using cryoSPARC (6). High-resolution reconstructions were obtained for both constructs, yielding structures of Spy-ApoF at 1.9 Å resolution and ALFA-ApoF at 1.98 Å resolution, and B-factors below 60 Å^2^ for both variants. A summary of the cryo-EM analysis of both Apo-ferritin constructs is shown in Figure 2. The reconstructed maps display well-resolved side chains and clear density for most residues. In several aromatic residues, the central hole of the ring system is visible, consistent with the high resolution of the reconstructions. Although SpyCatcher3 and the ALFA nanobody are bound to the ferritin subunits, the capture proteins were barely visible in the two-dimensional class averages and were not resolved in the final reconstructions due to averaging imposed by the octahedral symmetry of the ferritin particle.

**Figure 2:**
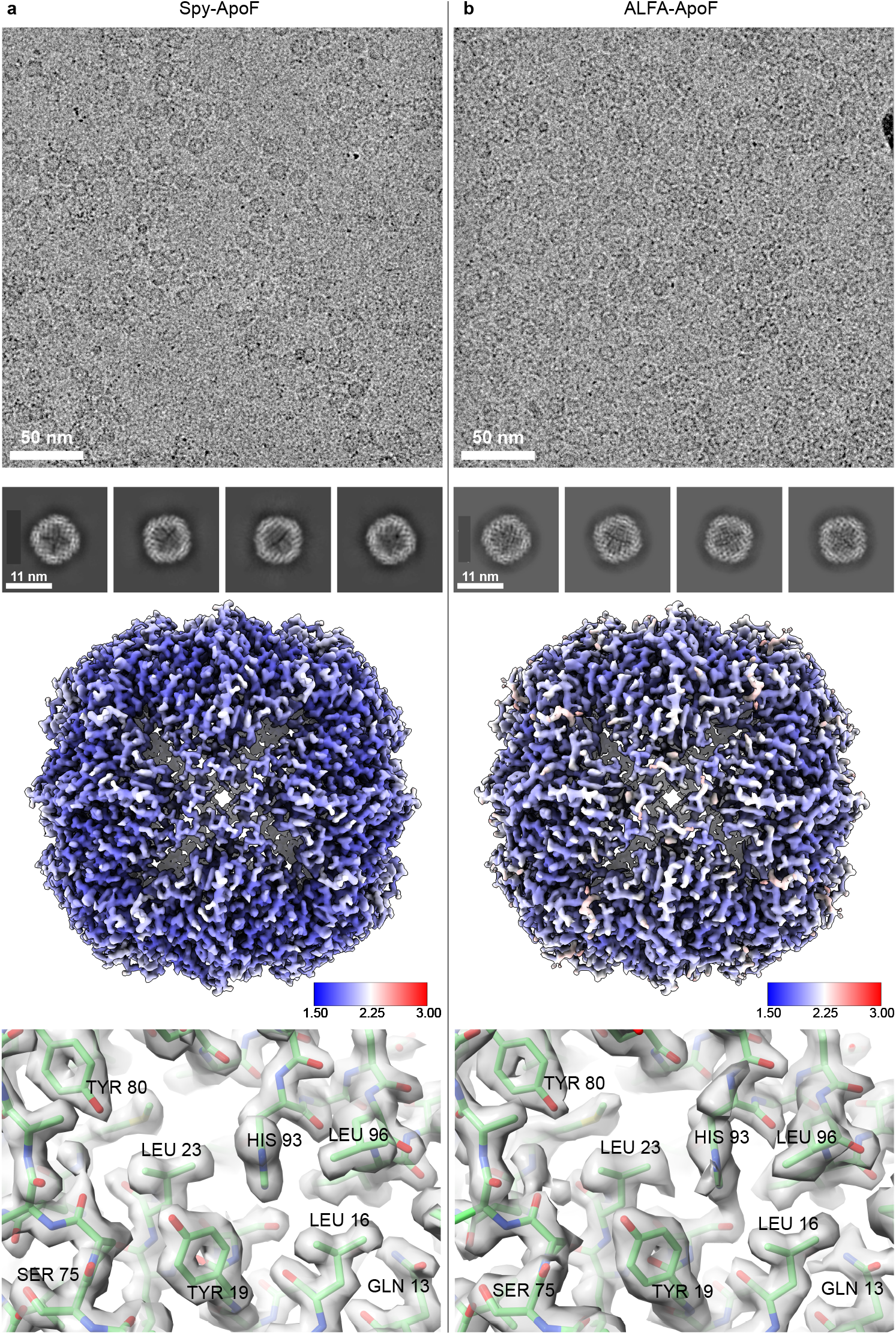
High-resolution Cryo-EM reconstructions of Apo-ferritin isolated from cell lysate using Spy- and ALFA-tags. **a)** Spy-ApoF (PDB: 28YP) and **b)** ALFA-ApoF (PDB: 28YQ). Representative micrographs and selected 2D-class averages are shown. Reconstructed maps are colored by local resolution (Å), and a selected region is shown with the fitted atomic model. Both structures could be refined to high resolution, with Spy-ApoF reaching 1.9 Å, ALFA-ApoF reaching 1.98 Å resolution and B-factors of 56.6 Å^*2*^ and 58.4 Å^*2*^, respectively, indicating high-quality grid preparation.

Atomic models were built starting from a previously determined ferritin structure (26) (PDB: 1EUM). The model was fitted into the reconstructed density and refined using Phenix (29) and Coot (30). Details of the processing workflow and model validation statistics are provided in Supplementary Information SI-6.

The resolution and overall map quality of the ALFA-ApoF reconstruction was slightly lower than those of Spy-ApoF, despite nearly twice as many particles contributing to the ALFA-ApoF reconstruction. We therefore examined whether differences in particle orientation distributions could explain this observation. In cryoSPARC, orientation bias can be quantified using the cFAR metric, which represents the ratio between the minimum and maximum weighted areas under the conical FSC curves and ranges from 0 to 1, with values closer to 1 indicating a more uniform angular distribution.

The Spy-ApoF reconstruction exhibited a cFAR value of 0.71, whereas ALFA-ApoF showed a significantly lower value of 0.37, indicating a stronger orientation bias. One possible explanation for this difference is the physicochemical distribution of hydrophilic and hydrophobic residues within the affinity tags. The ALFA-tag contains a more pronounced segregation of hydrophilic and hydrophobic surfaces, which may promote preferential adsorption of the fusion protein to the air-water interface during grid preparation and thereby restrict particle orientations. In contrast, hydrophilic and hydrophobic residues are more evenly distributed in SpyTag3, which may reduce such interactions and result in a more uniform orientation distribution. Notably, both capture proteins, the ALFA-nanobody and SpyCatcher3, display predominantly hydrophilic surfaces with comparable charge distributions, suggesting that the orientation bias arises from the small peptide tags themselves rather than from the bound capture reagents (see Supplementary Information SI-5).

### 2.4 Isolation of Additional Protein Targets from Cell Lysates

To further demonstrate the versatility of the microfluidic isolation workflow, we applied it to two additional proteins: β-galactosidase from *E. coli* and VgrG1 from *Pseudomonas aeruginosa*. Both proteins were expressed in *E. coli*, and clarified lysates were used directly for microfluidic isolation following the same workflow as described for Apo-ferritin.

For each target protein, constructs containing either SpyTag3 or ALFA-tag were prepared. All four variants could be successfully isolated using the cryoWriter (Supplementary Information SI-3B). For high-resolution cryo-EM analysis, one construct of each protein was selected. VgrG1-Spy was chosen due to its lower expression levels, where the covalent SpyTag3/SpyCatcher3 bond allows more stringent washing without losing bound protein. For β-galactosidase, the ALFA-tag construct was selected to provide an additional example of high-resolution reconstruction using ALFA-nanobody capture.

Cryo-EM data was collected on a FEI Titan Krios G4 and processed using cryoSPARC (6). Reconstructions were obtained at 2.63 Å resolution for VgrG1-Spy and 2.48 Å resolution for β-galactosidase-ALFA. An overview of the isolation workflow and representative reconstructions is shown in Figure 3. In two-dimensional class averages, faint densities corresponding to the bound capture proteins were visible: SpyCatcher3 adjacent to VgrG1 and the ALFA-nanobody bound to β-galactosidase. These densities were more apparent in the three-dimensional reconstructions (Supplementary Figs. S6-3 and S6-4). Note that the structures of the capture proteins themselves could not be resolved due to flexibility of the linker connecting the tag to the target protein, and, more importantly, they did not hinder the reconstruction of the target proteins.

**Figure 3:**
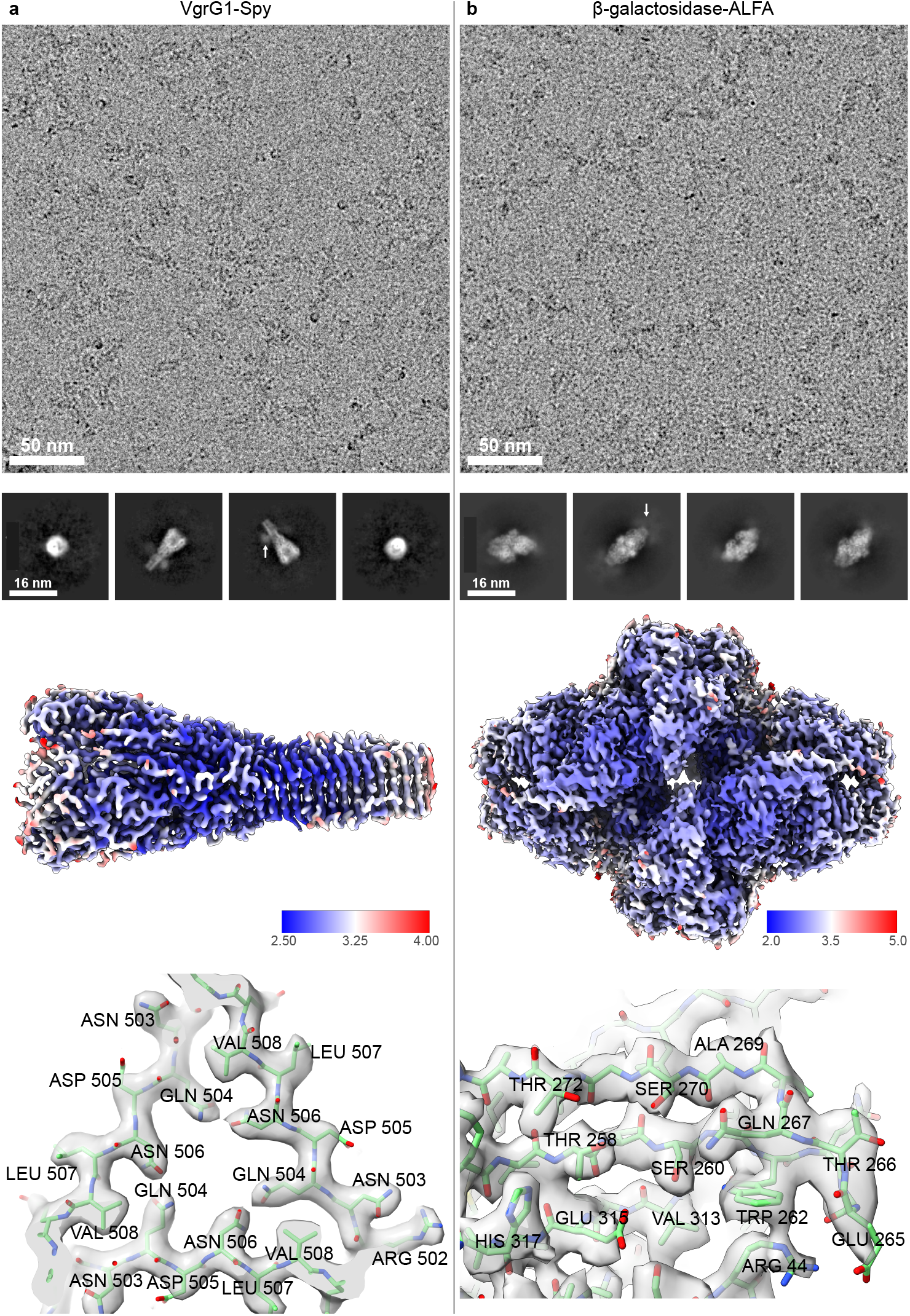
High-resolution cryo-EM reconstructions of VgrG1-Spy and β-galactosidase-ALFA isolated from cell lysate. **a)** VgrG1-Spy (PDB: 28YN) and **b)** β-galactosidase-ALFA (PDB: 28YO) isolated from 50 µL of cell lysate using the microfluidic workflow. Representative micrographs and selected 2D class averages are shown. The white arrows indicate density caused by the bound SpyCatcher or ALFA-nanobody. Reconstructed maps are colored by local resolution (Å), and a selected region is shown with the fitted atomic model. The reconstruction of VgrG1-Spy reached 2.63 Å and β-galactosidase-ALFA reached 2.48 Å resolution, with B-factors of 41 Å^*2*^ and 62.2 Å^*2*^, respectively, showing high-quality grid preparation.

Atomic models of VgrG1-Spy and β-galactosidase-ALFA were generated by fitting AlphaFold3 predictions into the reconstructed volumes by rigid-body fitting in ChimeraX, followed by refinement using Phenix and Coot. The resulting models show well-resolved secondary structure elements and side-chain densities consistent with the reported resolutions. Additional processing details and model validation statistics are provided in Supplementary Information SI-6.

### 2.5 Isolation of Ferritin A from *In Vitro* Translation Reactions

*In vitro* translation (IVT) reactions typically operate at microliter-scale volumes with lower yields than cellular expression, making them a demanding test case for the microfluidic workflow. A typical microfluidic isolation requires only ~10 pmol of protein, an amount readily produced in commercial IVT systems. To evaluate compatibility with IVT-derived samples, we produced ALFA-ApoF using a cell-free expression system and applied the same microfluidic isolation workflow used for lysate-derived samples. Expression of the protein was confirmed by pull-down assays using ALFA-nanobody-functionalized beads followed by SDS-PAGE analysis (see Supplementary Information SI-7).

During initial microfluidic isolation experiments we observed that samples used immediately after translation contained only a small fraction of fully assembled Apo-ferritin particles, along with many smaller particles (4–5 nm). Allowing the IVT reaction to mature overnight at 4 °C substantially increased the proportion of assembled ferritin particles, suggesting that oligomer assembly continues after translation (see Supplementary Information SI-7).

Following this optimization, microfluidic isolation of ALFA-ApoF from IVT reactions yielded cryo-EM grids with sufficient particle density for high-resolution data collection. The reconstruction reached 2.04 Å resolution with a B-factor of 67.4 Å^2^, comparable to the values obtained from lysate-derived samples (Spy-ApoF: 56.6 Å^2^; ALFA-ApoF: 58.4 Å^2^) and indicating that IVT-derived material yields cryo-EM samples of similar quality. We attribute the slightly higher B-factor to a residual fraction of incompletely assembled Apo-ferritin particles in the IVT-derived sample.

The atomic model was built starting from the ALFA-ApoF structure obtained from the lysate-derived sample and refined against the IVT-derived density map. The resulting model closely matches the lysate-derived structure (RMSD = 0.69 Å). The processing workflow and model validation statistics are provided in Supplementary Information SI-6, and an overview of the isolation and cryo-EM analysis is shown in Figure 4.

**Figure 4:**
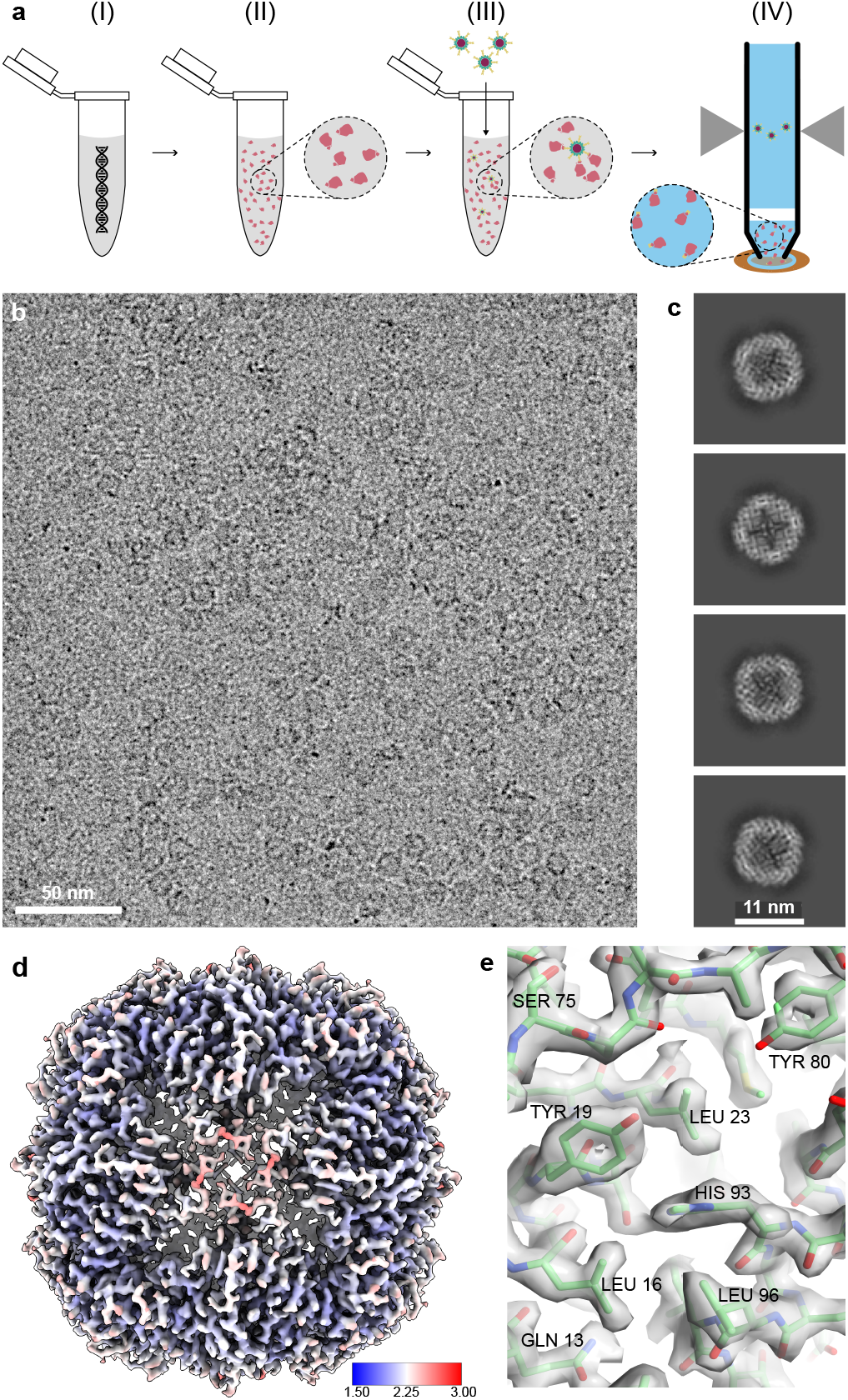
*In vitro* translation, isolation, and high-resolution cryo-EM reconstruction of ALFA-ApoF. **a)** Protein extraction was performed directly from a 50 µL *in vitro* translation (IVT) reaction. *(I)* The IVT reaction was initiated by adding plasmid DNA to the components of a commercial IVT kit and incubated for 3 h at 37 °C. *(II)* The reaction mixture was stored overnight at 4 °C to allow protein complex assembly. *(III)* Functionalized magnetic beads were added and incubated with the IVT sample for 1 h to capture the target protein. *(IV)* The microfluidic isolation workflow was performed as described in Figure 1. **b)** Representative cryo-EM micrograph of the microfluidic sample preparation. **c)** 2D class averages of ALFA-ApoF particles. **d)** Reconstructed 3D map colored by local resolution (Å). The gold-standard FSC (GSFSC) resolution is 2.04 Å, and the B-factor is 67.4 Å^*2*^. **e)** A representative region of the fitted atomic model (PDB: 28YR) shown within the reconstructed density.

These results demonstrate that microfluidic isolation enables direct structural analysis of proteins produced in cell-free systems, opening possibilities for rapid screening of protein variants and reaction conditions prior to cryo-EM analysis.

## 3 Discussion

The results presented here demonstrate that microfluidic protein isolation using affinity tags, combined with blotting-free cryo-EM grid preparation, enables high-resolution structural analysis directly from complex biological mixtures. We describe a microfluidic workflow, associated automation, and protocols for the isolation of tagged proteins from minimal amounts of starting material, extending our group’s previous work (17, 18).

Using our isolation approach (Figure 1), we reduce the required starting material by more than three orders of magnitude and shorten the workflow approximately 3-fold compared to conventional purification from *E. coli* overexpression cultures. Coupling the approach to *in vitro* translation reactions reduces the combined expression and purification timeline by up to 10-fold relative to conventional cell-based workflows.

Conventional chromatographic workflows typically require milliliter-scale input and multiple sequential steps that progressively dilute the target protein. This can destabilize weakly interacting complexes, reduce recovery of low-abundance species, and impose mechanical and chemical stress through repeated matrix interactions and buffer exchanges. Recent efforts have miniaturized chromatographic purification for cryo-EM sample preparation (35); our approach takes a fundamentally different and complementary route by combining batch capture with direct, blotting-free grid preparation.

Specifically, it relies on batch capture followed by photoelution within nanoliter-scale volumes and is directly integrated with automated cryo-EM grid preparation. Proteins are maintained in highly confined volumes throughout the workflow, thereby avoiding dilution while concentrating the material through extreme volume reduction (~50 µL input to ~25 nL output; >1000-fold volume compression). The batch format enables gentle removal of non-specifically bound contaminants, while photoelution provides a second, orthogonal purification step without requiring buffer exchange. Finally, the eluted protein is transferred directly onto the cryo-EM grid through the same nozzle used for isolation, eliminating intermediate handling steps between purification and vitrification.

The combination of bead trapping, photoelution, microfluidics, and associated robotics enables a range of adaptable isolation workflows that can be tailored to the starting material, input amount, target protein, and underlying biomedical question. In the present work, we focus on a workflow designed for high-throughput applications, in which bead binding is performed off-line in standard tubes (~50 µL volume; Figure 1, Figure 4). This configuration allows multiple protein preparations to be carried out in parallel and is compatible with standard liquid-handling automation, while the microfluidic platform is reserved for the final purification and grid preparation steps. After bead binding, the beads are concentrated to a final volume of ~8 µL, sufficient for more than 16 microfluidic isolations (100–500 nL of bead suspension per isolation), with each isolation taking ~15 min and yielding up to three cryo-EM grids or, alternatively, two cryo-EM grids and one negative-stain grid. These throughput characteristics are particularly advantageous when combined with *in vitro* translation systems, where substantial upconcentration is required and rapid screening of variants benefits from parallel preparation. For rare or limited biological material, an alternative, fully integrated in-line workflow is more appropriate. Building on our previous work targeting native, untagged protein complexes via antibody-based capture (18), we have applied this configuration to minimal cell volumes and patient-derived samples (manuscript in preparation), demonstrating that the platform can also isolate untagged endogenous proteins from limited material. The two configurations are complementary: tag-based parallel capture for highthroughput screening of recombinant or cell-free expressed targets, and in-line antibody-based capture for endogenous proteins from precious samples.

In this work, we compared several short affinity tags (<20 amino acids) within our microfluidic isolation workflow (Table 1) and identified two particularly robust and reproducible strategies: the ALFA-tag and the covalent SpyTag3/SpyCatcher3 system. In our batch isolation workflow, both the nanobody and SpyCatcher3 remain associated with the target protein after photoelution and co-elute with the complex; however, this does not interfere with downstream cryo-EM analysis, as both binders are small and predominantly hydrophilic. The ALFA-tag relies on high-affinity binding to an ALFA-specific nanobody, providing a reversible interaction between the target protein and the capture reagent. In contrast, the SpyTag3/SpyCatcher3 system forms a covalent bond, resulting in irreversible immobilization that is particularly advantageous for low-abundance targets, where stringent washing without target loss is essential. The capture reaction itself is rapid and proceeds efficiently at nanomolar concentrations (20). Importantly, both systems offer well-defined functionalization sites, allowing reliable coupling to photocleavable linkers without compromising binding performance. As a result, both tagging strategies enabled efficient isolation from lysates and *in vitro* translation reactions while remaining fully compatible with high-resolution cryo-EM.

A direct comparison of performance does not reveal a universally superior system, as the optimal choice depends on the target protein and the specific experimental context. In subsection 2.3 we observed higher recovery yields with ApoFerritin-ALFA, whereas SpyTag constructs produced grids of higher overall quality. The improved grid quality likely reflects two contributions: the covalent capture allows more stringent washing without target loss, reducing contamination; and the more uniform hydrophilic/hydrophobic distribution on SpyTag3 results in less preferential particle orientation at the air-water interface (subsection 2.3, SI-5). These observations highlight that tag selection is not merely a biochemical consideration but can directly impact cryo-EM sample quality through its inherent influence on interfacial and physicochemical behavior. While capture efficiencies are difficult to compare directly due to the fundamentally different binding mechanisms, both systems operate effectively in the nanomolar regime: the ALFA-nanobody through high-affinity equilibrium binding, and the SpyTag3/SpyCatcher3 system through rapid covalent bond formation. In practice, ALFA-tag capture is well-suited to abundant targets where high yield matters, while SpyTag3/SpyCatcher3 is preferable for low-abundance targets where wash stringency and grid quality dominate over absolute yield.

Compatibility with in vitro translation (IVT) systems is a key advantage of the workflow. As proof-of-concept, ALFA-tagged ApoF produced by IVT was successfully isolated and yielded a high-resolution reconstruction with favorable B-factor, demonstrating that microfluidic isolation provides sufficient material of sufficient quality for cryo-EM analysis from cell-free reactions. Combined with the parallel workflow described above, this opens opportunities for rapid and flexible protein production, enabling high-throughput structural screening of protein variants or reaction conditions. This may be particularly valuable for proteins difficult to express in cells, including toxic targets, those requiring unnatural amino acids, and those benefiting from defined or systematically varied reaction conditions (36–38).

Beyond isolation chemistry, the deposition step critically shapes data quality. A minimal amount of eluted protein is required to offset adsorption to the carbon support, which would otherwise reduce particle density. In our workflow, ~20 nL of sample is dispensed onto the grid and excess liquid is re-aspirated, leaving a thin layer that concentrates proteins within the holes. This deposition-reaspiration strategy yields particle densities suitable for high-resolution cryo-EM while keeping consumption low, enabling up to three grids from a single microfluidic isolation. Other grid-writing approaches, including microfluidic dispensing (39) and pin-based deposition (40), use smaller volumes but require higher protein concentrations (typically 5–15 mg mL^−1^) to achieve comparable densities. The optimal deposition strategy thus depends on whether the upstream workflow provides dilute or concentrated samples; deposition-reaspiration is particularly well suited for the dilute eluate generated by microfluidic capture from minimal starting material.

The main limitation of the approach arises from the very small amounts of material processed.Unlike classical purification workflows, where intermediate fractions can be sampled and analyzed (e.g., by Western blotting or mass spectrometry), such monitoring is largely impractical here: the protein is immobilized on beads through most of the workflow, and the eluate volumes are too small to spare aliquots for routine parallel analysis. Several strategies partially mitigate this. First, optical readouts enable real-time tracking of bead enrichment and photoelution release whenever the target protein (e.g., a GFP fusion) or the capture reagent carries a fluorescent label. Second, the cryoWriter capillary can dispense eluate aliquots for orthogonal downstream analyses such as reverse-phase protein arrays (41) or mass spectrometry. Finally, multiple grids are typically prepared from a single isolation, and negative-stain EM of one of them provides a rapid quality control step that distinguishes failures in protein isolation (e.g., damage or loss) from those occurring during cryo-EM grid preparation.

Future developments will focus on expanding compatible capture strategies and addressing more challenging targets. One promising direction is the use of endogenously tagged proteins generated by CRISPR/Cas-based genome editing (42, 43). In this context, the integrated fluorescence microscope of the cryoWriter could be combined with microfluidic isolation to identify and process GFP-tagged proteins directly from few or even single cells. As a proof of concept,we successfully implemented GFP-based capture within the microfluidic workflow (Supplementary Information SI-3C), although no structure was obtained, likely due to target flexibility or instability. More broadly, application of the workflow to detergent-solubilized membrane-protein constructs has been demonstrated at the level of capture and grid preparation; obtaining high-resolution structures of such targets, where construct stability under cryo-EM conditions is the primary challenge, is the subject of ongoing work.

In summary, microfluidic protein isolation combined with blotting-free cryo-EM grid preparation enables high-resolution structural analysis directly from complex biological mixtures using minimal starting material. By integrating batch capture, specific photoelution, and automated deposition, the workflow produces cryo-EM samples suitable for high-resolution structure determination. Continued development of such approaches is expected to enable structural analysis of increasingly challenging systems and to support integrated workflows at the interface of biochemistry, cell biology, and structural biology.

## 4 Methods

### 4.1 Cloning and Protein Expression

#### 4.1.1 *E.Coli* FtnA

A plasmid containing the sequence for *E.Coli* Ferritin A (FtnA) was a generous gift from Dr. Babatunde Ekundayo and Prof. Henning Stahlberg at EPFL. FtnA was cloned into a pETDuet1-vector by GenScript along with a N-terminal 6x His-tag followed by SpyTag3 (20). A similar construct was also made with an ALFA-tag (19) instead of SpyTag3. We call these constructs Spy-ApoF and ALFA-ApoF, respectively. The plasmids were used to transform *E.Coli* BL21 DE3 competent cells (ThermoFisher, EC0114). These cells were grown at 37 °C for 16 h on agar-plates containing 100 µg mL^−1^ ampicillin.

For protein expression, one colony was selected and the cells were grown overnight in 2YT-medium (supplemented with 100 µg mL^−1^ ampicillin, 100 mmol dm^−3^ phosphate buffer and 1 % w/v D-(+)-Glucose (>99.5 %; Sigma, G8270)) at 37 °C and 110 RPM. 20 mL of this pre-culture were used to start the main culture (0.8 L); protein expression was induced at OD_600_ = 0.7 using 0.4 mmol dm^−3^ IPTG (>99 %; LubioScience, LU5004) and the culture was subsequently incubated for 4 h at 37 °C and 110 RPM.

The grown cells were harvested by centrifugation at 5000 ×g for 10 min. The cell-pellet was resuspended in 20 mL resuspension buffer (50 mmol dm^−3^ HEPES, 300 mmol dm^−3^ NaCl, 10 mmol dm^−3^ MgCl_2_, 10 % Glycerol, pH7.5) supplemented with 1 pill of cOmplete™ Mini, EDTA-free Protease-Inhibitor-Cocktail (Roche, 11836170001) and 250 µL DNase I (2 mg mL^−1^; IWT Reagents, A3778). The cells were lysed by sonication and the lysate was cleared by ultracentrifugation at 41 000 RPM for 1 h at 6 °C. The supernatant was aliquoted, flash-frozen in liquid nitrogen and stored at −80 °C.

#### 4.1.2 *E.Coli* β-galactosidase

The plasmid encoding *E.Coli* β-galactosidase (pDR111-mannose-Bgal NLS from Christopher Contag (21); Addgene plasmid 188398) was purchased from AddGene. The sequence encoding β-galactosidase was cloned in a pET-15b vector by GenScript along with a C-terminal ALFA-tag (19) or SpyTag3 (20). We call these constructs β-galactosidase-ALFA and β-galactosidase-Spy, respectively.

The expression protocol was the same as described for *E.Coli* FtnA, except that the resuspension buffer was 50 mmol dm^−3^ HEPES, 200 mmol dm^−3^ NaCl, 1 mmol dm^−3^ MgCl_2_, pH8.

#### 4.1.3 *Pseudomonas aeruginosa* VgrG1

The plasmid encoding *Pseudomonas aeruginosa* VgrG1 was a generous gift from Dr. Mitchell Brüderlin, Prof. Roderick Lim and Prof. Marek Basler from the University of Basel. VgrG1 was cloned into a pET-Duet vector by GenScript along with a C-terminal SpyTag3 (20) or ALFA-tag (19). We call these constructs VgrG1-Spy and VgrG1-ALFA, respectively. The expression protocol was the same as described for *E.Coli* FtnA, except that the resuspension buffer was 20 mmol dm^−3^ Tris, 150 mmol dm^−3^ NaCl, 5 mmol dm^−3^ MgCl_2_, pH7.8.

#### 4.1.4 *In Vitro* Translation (IVT) of *E.Coli* FtnA

ALFA-FtnA was produced *in vitro* using a PURExpress *In Vitro* Protein Synthesis Kit (New England Biolabs, NEB-E6800). The kit was a kind gift from Mette Roesgaard from the group of Jan Pieter Abrahams of the University of Basel. Briefly, 20 µL solution A and 15 µL solution B were mixed together, then 2 µL of RNase inhibitor (applied biosystems, N8080119) and 13 µL (i.e. 294 ng) plasmid DNA were added, leading to a total sample volume of 50 µL. This mixture was incubated on a shaker for 3 h at 37 °C and 500 RPM. The IVT-sample was stored overnight at 4 °C and used the following day for protein isolation. 1 µL DNase I (2 mg mL^−1^; IWT Reagents, A3778) was added to the IVT-sample on the day of the protein isolation to digest the remaining plasmid DNA.

#### 4.1.5 Cold-Shock DEAD-box Protein A (CsdA)

CsdA was expressed as described elsewhere (22) and the cleared lysate containing CsdA with a C-terminal maltose-binding protein (MBP) tag and a N-terminal GFP-tag was a kind gift from Michelle Gut (University of Basel, Hondele Lab). The resuspension buffer for CsdA was 25 mmol dm^−3^ Tris, 150 mmol dm^−3^ NaCl, 2 mmol dm^−3^ MgCl_2_, 0.5 mmol dm^−3^, β-mercaptoethanol, 10 % glycerol, pH(7.5). The lysate was aliquoted, flash-frozen in liquid nitrogen and stored at −80 °C.

### 4.2 Bio-layer Interferometry Measurements

Bio-layer interferometry (BLI) measurements were made using a BLItz− system (Pall Life Sciences, FortéBio). Streptavidin-coated biosensors were functionalized with bioinylated ALFA-nanobodies (2.8 µmol dm^−3^ in PBS pH7.5).

Briefly, a 30 s baseline was recorded, then the nanobodies were loaded onto the biosensor for 300 s and lastly unbound nanobodies were allowed to dissociate for 60 s. The binding assay was conducted with the functionalized biosensor by recording a 30 s baseline followed by 180 s of association and 180 s of dissociation of the target protein. The assay was carried out for different concentrations of ALFA-ApoF (12.5, 50 and 200 nmol dm^−3^ in 50 mmol dm^−3^ HEPES, 150 mmol dm^−3^ NaCl, pH7.5). The measured data was analysed using the BLItz− Pro software in order to determine the dissociation constant *K*_*D*_. Similar assays were performed for GFP-tagged CsdA and His-tagged GFP in order to determine the binding affinity of the anti-GFP nanobody (ChromoTek, gt-250) and the anti-His Fabs (Cytiva, 27-4710-01), respectively (see SI-4).

### 4.3 Microscale Thermophoresis Measurements

Microscale thermophoresis (MST) measurements were made using a Monolith NT.115 system (NanoTemper Technologies). The binding assay was performed by titrating the anti-His RNA aptamers (23) against 20 nmol dm^−3^ His-tagged GFP (Buffer: 50 mmol dm^−3^ HEPES, 100 mmol dm^−3^ NaCl, 0.1 mmol dm^−3^ magnesium acetate, 0.01 % Tween-20, pH7.2). Data was recorded for 16 different dilutions and the measured data was analysed using the MO.Affinity Analysis software (NanoTemper Technologies) in order to determine the dissociation constant *K*_*D*_ (see SI-4).

### 4.4 Pull-Down using ALFA-Nanobodies

The pull-down using ALFA-nanobodies was done using 10 µL of streptavidin-coated magnetic beads (ThermoFisher, T1 DynaBeads, 65601), functionalized with 45 pmol ALFA-nanobodies. The beads were incubated with 100 µL of cell lysate for 1 h at room temperature. Then, the beads were pulled down using a magnet and the supernatant was removed. The beads were washed (5 × 50 µL) with PBS and resuspended in 10 µL PBS. An equal amount of 2x SDS-loading buffer was added and the samples were heated to 95 °C for 5 min. 10 µL were loaded onto a NuPAGE− Bis-Tris 4–12 % polyacrylamide gel (ThermoFisher, NP0321BOX) and the gel was run at 150 V for 50 min. 6 µL Precision Plus Protein™ Dual Color Standard (BioRad, 1610374) were used as a molecular weight marker on the SDS-PAGE gels. The gels were stained using Quick Blue Protein Stain (LubioScience, LU001000).

### 4.5 Fluorescence Imaging of SDS-PAGE Gel

SpyCatcher3-CYS was fluorescently labelled using a 10 × molar excess of CF− 647 maleimide dye (Biotium, 92027). The excess dye was removed using a Zeba Micro-Spin Desalting Column (ThermoFisher, 89877, 7k MWCO). 10 µL of cell lysate were incubated with 125 pmol of labelled SpyCatcher for 1 h at room temperature. Then, an equal amount of 2x SDS-loading buffer was added to the samples and they were heated to 95 °C for 5 min. 10 µL were loaded onto a NuPAGE− Bis-Tris 4–12 % polyacrylamide gel (ThermoFisher, NP0321BOX) and the gel was run at 150 V for 50 min. The unstained gel was imaged with an infra-red imager (LI-COR Odyssey Fc, LICORbio) using the 700 nm channel and integrating for 5 min. The Image Studio− software (LICORbio) was used to analyse and export the images. After the fluorescent imaging, the gel was stained using Quick Blue Protein Stain (LubioScience, LU001000). 6 µL Precision Plus Protein− Dual Color Standard (BioRad, 1610374) were used as a molecular weight marker on the SDS-PAGE gels. This protocol was adapted from (24).

### 4.6 Biotinylation via Thiol-groups

Lyophilized ALFA-nanobodies (19) with a C-terminal cysteine (NanoTag Biotechnologies, N1505) were resuspended in PBS to a concentration of 1 mg mL^−1^. In a first step, the EZ-link™ maleimide-DBCO crosslinker (ThermoFisher, C20041) was bound to the nanobodies using a 10 × molar excess. After 2 h at room temperature, the excess crosslinker was removed using a Zeba Micro-Spin Desalting Column (ThermoFisher, 89882, 7k MWCO). In a second step, a 10 × molar excess of PC-Biotin-PEG3-Azide crosslinker (BroadPharm, BP-22676) was added and incubated for 16 h in the dark at 4 °C. The excess crosslinker was removed using a Spin Column, as before. The biotinylated ALFA-nanobodies were stored protected from light at 4 °C. If stored and handled properly, they remained stable for at least 6 months.

SpyCatcher3-CYS (20) (BioRad, TZC025CYS) was incubated with 5 mmol dm^−3^ DTT for 30 min in order to break the disulfide bonds between cysteines. The DTT was removed using a Zeba Micro-Spin Desalting Column (ThermoFisher, 89882, 7k MWCO). The rest of the biotinylation protocol of SpyCatcher was the same as for the ALFA-nanobodies. Biotinylated SpyCatcher has also remained stable for at least 6 months.

### 4.7 Biotinylation via Primary Amines

Capture reagents may also be biotinylated via primary amines, which are present on the side chains of lysines and the N-terminus proteins. To do so, a crosslinker containing an NHS-ester group is used. In our experiments, we used PC Biotin-NHS ester (Vector Labs, CCT-1225-1) to biotinylate Tris-NTA-amine-crosslinker (biotechrabbit, BR1001101), Anti-His Antibody (Cytiva, 27-4710-01), ANTI-FLAG® M2 antibody (Sigma, F1804) and ChromoTek anti-GFP recombinant VHH (ChromoTek, gt-250).

In short, the capture reagent was incubated with a 10 × molar excess of the NHS-ester crosslinker for 1 h at room temperature. The excess crosslinker was removed using a Zeba Micro-Spin Desalting Column (ThermoFisher) with the appropriate molecular weight cutoff, except for the Tris-NTA-amine-crosslinker, which was bound directly to the magnetic beads. The biotinylated capture reagents were stored protected from light at 4 °C.

### 4.8 Protein Isolation Workflow

Streptavidin-coated magnetic beads (ThermoFisher, T1 DynaBeads, 65601) were functionalized with 4 pmol of biotinylated binder per 1 µL of beads. A typical experiment requires 2–4 µL of beads. The functionalized beads were washed 3 × in 50 µL PBS to remove unbound nanobodies or SpyCatcher. The functionalized beads were incubated with 10–100 µL cell lysate or 50 µL IVT-sample for 1 h on an overhead shaker at 4 °C.

After incubation, the beads were collected at the bottom of the tube using a magnet. The supernatant was carefully removed and the beads were washed (5 × 50 µL) with the appropriate buffer. The Apoferritin-sample was washed using 50 mmol dm^−3^ Tris, 50 mmol dm^−3^ NaCl, 1 mmol dm^−3^ EDTA, pH7.5, the β-galactosidase sample with 25 mmol dm^−3^ Tris, 50 mmol dm^−3^ NaCl, 2 mmol dm^−3^ MgCl_2_, 1 mmol dm^−3^ TCEP, pH8 and the VgrG1 sample with 20 mmol dm^−3^ Tris, 150 mmol dm^−3^ NaCl, pH8. After washing, the beads were resuspended in 8 µL buffer.

30 nL of air were aspirated in the microcapillary and then 100–500 nL beads were taken up and captured in the magnetic trap. This plug of beads was then moved to the tip of the capillary and the air-bubble was also moved forwards such that the beads were enclosed in a volume of ~30 nL. This small volume was then moved back slightly in order to introduce an air-bubble in front of it, followed by 20 nL of buffer to avoid evaporation of the liquid around the beads. Next, the protein was eluted from the beads by applying UV-light (365 nm, 190 mW, Thorlabs; M365L2) for 15 min. After photo-cleaving, cryo- or negative stain grids were prepared. For cryo-grids, 17 nL of the eluate were deposited on a grid (Quantifoil, Cu300, R2/1), which was subsequently plunge-frozen in liquid ethane. The grid had been glow-discharged for 60 s in air plasma immediately before use. The grid environment was held at 80 % relative humidity using the ClimateJet (15), and the dew-point stage was set 3 °C below the dew point of the ClimateJet atmosphere. In steady state this brings the grid surface close to the local dew point, controling evaporation without inducing condensation. For negative stain grids, the sample can either be prepared classically; i.e. deposit the sample on the grid, wash it with pure water and stain it with 2 % uranyl acetate, or it can be prepared directly using the cryoWriter (25).

### 4.9 Data Collection

Cryo-EM data were collected on a Titan Krios G4 (Thermo Fisher Scientific) transmission electron microscope operated at 300 kV, equipped with a Selectris X energy filter set to 10 eV and a Falcon 4i direct electron detector operated in counting mode. Movies were recorded at a nominal magnification of 165 000 (pixel size of 0.73 Å) with a total exposure time of 3 s split into 60 frames. The defocus-range was set to 0.8–2 µm and the total electron dose was 40–45 e^−^/Å^2^, depending on the dataset. Automated data acquisition was performed using Smart EPU (Thermo Fisher Scientific).

### 4.10 Image Processing

Image processing was performed in cryoSPARC (6). Movies were motion- and CTF-corrected, and high-quality micrographs were retained for downstream processing. Particles were blob-picked on a random subset of micrographs, extracted, and 2D-classified to generate templates. Template-based picking was then run on the full dataset, followed by iterative 2D classification to remove poor-quality particles. An *ab initio* model was generated from the cleaned particle set and refined using homogeneous or non-uniform refinement with the appropriate symmetry and higher-order CTF corrections. Reference-based motion correction was applied, and a final refinement yielded the high-resolution reconstruction. Processing parameters and refinement statistics are reported in Supplementary Information SI-6.

### 4.11 Model Building

The model of FtnA was built based on the X-ray structure of Stillman *et al*. (26) (PDB-ID: 1EUM). Initial models for VgrG1 and β-galactosidase were generated with AlphaFold3 (7). All models were rigid-body fitted into the reconstructed densities in ChimeraX (27, 28), followed by real-space refinement in Phenix v.2.0 (29) and manual adjustments in Coot v.1.1.15 (30). Figures of models and densities were prepared in ChimeraX. Model validation statistics are reported in Supplementary Information SI-6.

## Acknowledgments

We thank the following Biozentrum facilities at the University of Basel for ongoing support and helpful discussions: the BioEM lab (Dr. M. Chami, Dr. C. F. Rodriguez, Dr. R. Ruedas), the Proteomics facility (Dr. A. Schmidt, Dr. D. Ritz), and the Biophysics facility (Dr. T. Sharpe, Dr.T. Mühlethaler). We acknowledge support from the Swiss Nanoscience Institute (ARGOVIA FuncEM project) and the Swiss National Science Foundation (projects 200021_162521 and 200020_192190) to T.B., as well as ROCHE ROADS funding (granted to T. B., M. L., and T. C.).

## Author Contributions

M.Z. and T.B. designed the research; T.C., R.T. and M.L. contributed to the study plan (ROCHE ROADS project) and enabled various test systems; M.Z. performed experiments with His-, FLAG-, GFP-, ALFA- and SpyTag3-tagged proteins, developed and performed quality control experiments, collected and analyzed EM data, and built protein models; E.S. and L.R. performed experiments with His- and GFP-tagged proteins and contributed to figures and discussions; T.B. supervised the project; M.Z. and T.B. wrote the manuscript with input from all authors.

