## Supplementary Information for "Affinity-tag-based microfluidic protein isolation enables high-resolution Cryo-EM from minimal starting material"

**Bibliography** **36**

### 1 Supplementary Information

#### 1.1 SI-1: Experimental System and Constructs

Additional information on protein constructs, plasmids, tagging strategies, and functionalization chemistry used for microfluidic capture.

##### 1.1.1 SI-1A Used Crosslinking Chemistry

The microfluidic isolation workflow relies on superparamagnetic particles that can be immobilized in the capillary using strong magnetic field gradients. The workflow contains two key steps:

- Specific binding of a capture reagent to the affinity tag of the target protein.
- Specific release of the captured protein by photocleavage.

In all cases, capture reagents were immobilized on streptavidin-coated magnetic beads via biotinylated photocleavable linkers, enabling release of the capture reagent-protein complex upon UV illumination. These linkers allow stable immobilization of the capture molecules during the binding and washing steps while enabling controlled release of the captured protein after the washing procedure.

The photocleavable crosslinkers used in this work and their functional groups are illustrated in Figure 1. Detailed information on the reagents and functionalization procedures is provided in the Materials and Methods section.

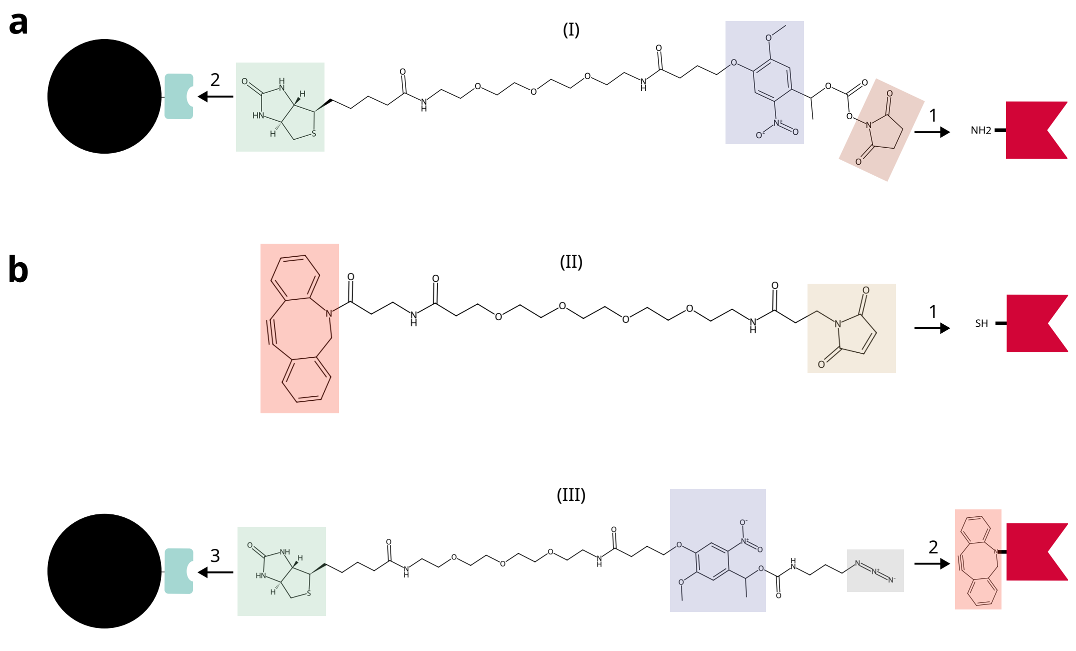

**Figure 1: Schematic overview of binder biotinylation strategies: a)** Primary amine coupling. The photocleavable (violet) linker (I) is coupled to primary amines on the capture reagent via its NHS-ester group (dark brown). The resulting biotinylated (green) binder is subsequently immobilized on streptavidin-coated magnetic beads. **b)** Thiol-directed coupling. The binder is first functionalized via maleimide (light brown) chemistry with a linker containing a DBCO group (red) (II). A biotin linker bearing an azide group (gray) (III) is then attached via strain-promoted azide-alkyne cycloaddition (click chemistry) between the azide and DBCO groups. The resulting biotinylated binder is finally immobilized on streptavidin-coated magnetic beads.

##### 1.1.2 SI-1B Protein Constructs and Plasmids

The plasmids encoding the constructs used in this study are summarized in Figure 2. To evaluate the general applicability of the microfluidic isolation workflow, we designed constructs representing proteins of different size and symmetry, including the well-established cryo-EM benchmark protein ferritin A (FtnA), the large tetrameric enzyme $\beta$-galactosidase, and the elongated trimeric protein VgrG1. Each target protein was fused to either SpyTag3 or the ALFA-tag to enable isolation using the corresponding capture reagents and for FtnA an additional His-tag is incorporated. The amino acid sequences of all constructs are listed below.

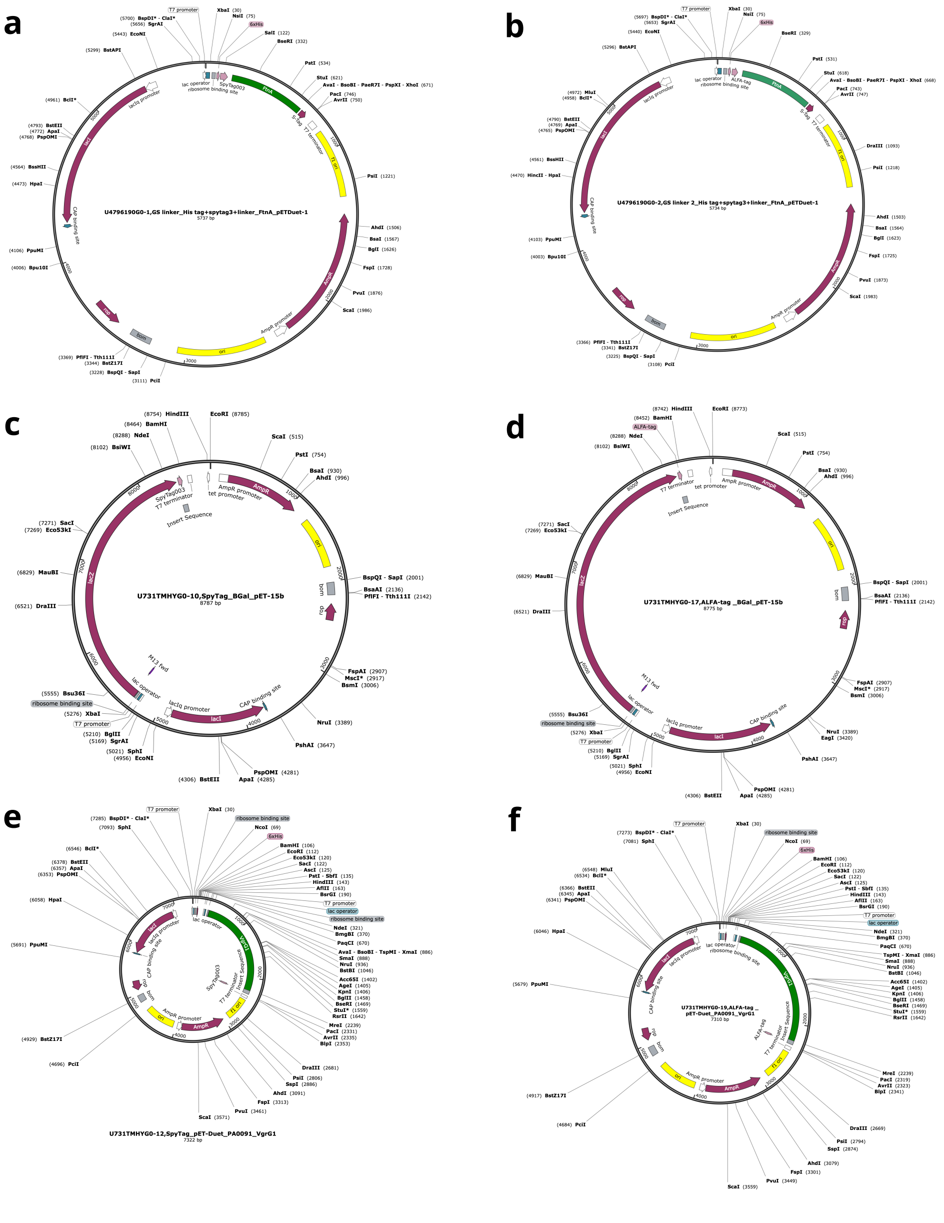

**Figure 2: Plasmids of all constructs: a)** pETDuet-1 vector containing Ferritin A with N-terminal His6-SpyTag3. **b)** pETDuet-1 vector containing Ferritin A with N-terminal His6-ALFA-tag. **c)** pET-15b vector encoding $\beta$-galactosidase with a C-terminal SpyTag3. **d)** pET-15b vector encoding $\beta$-galactosidase with a C-terminal ALFA-tag. **e)** pETDuet-1 vector encoding VgrG1 with a C-terminal SpyTag3. **f)** pETDuet-1 vector encoding VgrG1 with a C-terminal ALFA-tag.

- Ferritin A with a N-terminally fused His$_6$-tag, a SpyTag3 and a GS linker sequence: MHHHHHHGSRGVPHIVMVDAYKRYKGSLKPEMIEKLNEQMNLELYSSLLY QQMSAWCSYHTFEGAAAFLRRHAQEEMTHMQRLFDYLTDTGNLPRINTV ESPFAEYSSLDELFQETYKHEQLITQKINELAHAAMTNQDYPTFNFLQWYV SEQHEEEKLFKSIIDKLSLAGKSGEGLYFIDKELSTLDTQN

- Ferritin A with a N-terminally fused His$_6$-tag, an ALFA-tag and a GSGSGSGSGS linker sequence: MHHHHHHGSPSRLEEELRRRLTEPGSGSGSGSGSLKPEMIE KLNEQMNLELYSSLLYQQMSAWCSYHTFEGAAAFLRRHAQEEMTHMQRL FDYLTDTGNLPRINTVESPFAEYSSLDELFQETYKHEQLITQKINELAHAAM TNQDYPTFNFLQWYVSEQHEEEKLFKSIIDKLSLAGKSGEGLYFIDKELSTL DTQN

- $\beta$-galactosidase with a C-terminally fused SpyTag3 and a GSGESGSG linker sequence: MTMITDSLAVVLQRRDWENPGVTQLNRLAAHPPFASWRNSEEAR TDRPSQQLRSLNGEWRFAWFPAPEAVPESWLECDLPEADTVVVPSNWQM HGYDAPIYTNVTYPITVNPPFVPTENPTGCYSLTFNVDESWLQEGQTRIIF DGVNSAFHLWCNGRWVGYGQDSRLPSEFDLSAFLRAGENRLAVMVLRWS DGSYLEDQDMWRMSGIFRDVSLLHKPTTQISDFHVATRFNDDFSRAVLEA EVQMCGELRDYLRVTVSLWQGETQVASGTAPFGGEIIDERGGYADRVTLR LNVENPKLWSAEIPNLYRAVVELHTADGTLIEAEACDVGFREVRIENGLLL LNGKPLLIRGVNRHEHHPLHGQVMDEQTMVQDILLMKQNNFNAVRCSHY PNHPLWYTLCDRYGLYVVDEANIETHGMVPMNRLTDDPRWLPAMSERVT RMVQRDRNHPSVIIWSLGNESGHGANHDALYRWIKSVDPSRPVQYEGGGA DTTATDIICPMYARVDEDQPFPAVPKWSIKKWLSLPGETRPLILCEYAHAM GNSLGGFAKYWQAFRQYPRLQGGFVWDWVDQSLIKYDENGNPWSAYGG DFGDTPNDRQFCMNGLVFADRTPHPALTEAKHQQQFFQFRLSGQTIEVTS EYLFRHSDNELLHWMVALDGKPLASGEVPLDVAPQGKQLIELPELPQPESA GQLWLTVRVVQPNATAWSEAGHISAWQQWRLAENLSVTLPAASHAIPHLT TSEMDFCIELGNKRWQFNRQSGFLSQMWIGDKKQLLTPLRDQFTRAPLDN DIGVSEATRIDPNAWVERWKAAGHYQAEAALLQCTADTLADAVLITTAHA WQHQGKTLFISRKTYRIDGSGQMAITVDVEVASDTPHPARIGLNCQLAQV AERVNWLGLGPQENYPDRLTAACFDRWDLPLSDMYTPYVFPSENGLRCG TRELNYGPHQWRGDFQFNISRYSQQQLMETSHRHLLHAEEGTWLNIDGFH MGIGGDDSWSPSVSAELQLSAGRYHYQLVWCQKGSGESGSGRGVPHIVMV DAYKRYK

- $\beta$-galactosidase with a C-terminally fused ALFA-tag and a GSGSGS linker sequence: MTMITDSLAVVLQRRDWENPGVTQLNRLAAHPPFASWRNSEEAR TDRPSQQLRSLNGEWRFAWFPAPEAVPESWLECDLPEADTVVVPSNWQM HGYDAPIYTNVTYPITVNPPFVPTENPTGCYSLTFNVDESWLQEGQTRIIF DGVNSAFHLWCNGRWVGYGQDSRLPSEFDLSAFLRAGENRLAVMVLRWS DGSYLEDQDMWRMSGIFRDVSLLHKPTTQISDFHVATRFNDDFSRAVLEA EVQMCGELRDYLRVTVSLWQGETQVASGTAPFGGEIIDERGGYADRVTLR LNVENPKLWSAEIPNLYRAVVELHTADGTLIEAEACDVGFREVRIENGLLL LNGKPLLIRGVNRHEHHPLHGQVMDEQTMVQDILLMKQNNFNAVRCSHY PNHPLWYTLCDRYGLYVVDEANIETHGMVPMNRLTDDPRWLPAMSERVT RMVQRDRNHPSVIIWSLGNESGHGANHDALYRWIKSVDPSRPVQYEGGGA DTTATDIICPMYARVDEDQPFPAVPKWSIKKWLSLPGETRPLILCEYAHAM GNSLGGFAKYWQAFRQYPRLQGGFVWDWVDQSLIKYDENGNPWSAYGG

DFGDTPNDRQFCMNGLVFADRTPHPALTEAKHQQQFFQFRLSGQTIEVTS EYLFRHSDNELLHWMVALDGKPLASGEVPLDVAPQGKQLIELPELPQPESA GQLWLTVRVVQPNATAWSEAGHISAWQQWRLAENLSVTLPAASHAIPHLT TSEMDFCIELGNKRWQFNRQSGFLSQMWIGDKKQLLTPLRDQFTRAPLDN DIGVSEATRIDPNAWVERWKAAGHYQAEAALLQCTADTLADAVLITTAHA WQHQGKTLFISRKTYRIDGSGQMAITVDVEVASDTPHPARIGLNCQLAQV AERVNWLGLGPQENYPDRLTAACFDRWDLPLSDMYTPYVFPSENGLRCG TRELNYGPHQWRGDFQFNISRYSQQQLMETSHRHLLHAEEGTWLNIDGFH MGIGGDDSWSPSVSAELQLSAGRYHYQLVWCQKGSGSGSPSRLEEELRRRL TE

- VgrG1 with a C-terminally fused SpyTag3 and a GSGESGSG linker sequence: M QLTRLVQVDCPLGPDVLLLQRMEGREELGRLFAYELHLVSENPNLPLEQLL GKPMSLSLELPGGSRRFFHGIVARCSQVAGHGQFAGYQATLRPWPWLLTR TSDCRIFQNQSVPEIIKQVFRNLGFSDFEDALTRPYREWEYCVQYRETSFDF ISRLMEQEGIYYWFRHEQKRHILVLSDAYGAHRSPGGYASVPYYPPTLGHR ERDHFFDWQMAREVQPGSLTLNDYDFQRPGARLEVRSNIARPHAAADYPL YDYPGEYVQSQDGEQYARNRIEAIQAQHERVRLRGVVRGIGAGHLFRLSG YPRDDQNREYLVVGAEYRVVQELYETGSGGAGSQFESELDCIDASQSFRLL PQTPVPVVRGPQTAVVVGPKGEEIWTDQYGRVKVHFHWDRHDQSNENSS CWIRVSQAWAGKNWGSMQIPRIGQEVIVSFLEGDPDRPIITGRVYNAEQTV PYELPANATQSGMKSRSSKGGTPANFNEIRMEDKKGAEQLYIHAERNQDN LVENDASLSVGHDRNKSIGHDELARIGNNRTRAVKLNDTLLVGGAKSDSVT GTYLIEAGAQIRLVCGKSVVEFNADGTINISGSAFNLYASGNGNIDTGGRLD LNSGGASEVDAKGKGVQGTIDGQVQAMFPPPAKGGSGESGSGRGVPHIVM VDAYKRYK

- VgrG1 with a C-terminally fused ALFA-tag and a GSGSGS linker sequence: MQ LTRLVQVDCPLGPDVLLLQRMEGREELGRLFAYELHLVSENPNLPLEQLLG KPMSLSLELPGGSRRFFHGIVARCSQVAGHGQFAGYQATLRPWPWLLTRT SDCRIFQNQSVPEIIKQVFRNLGFSDFEDALTRPYREWEYCVQYRETSFDFI SRLMEQEGIYYWFRHEQKRHILVLSDAYGAHRSPGGYASVPYYPPTLGHR ERDHFFDWQMAREVQPGSLTLNDYDFQRPGARLEVRSNIARPHAAADYPL YDYPGEYVQSQDGEQYARNRIEAIQAQHERVRLRGVVRGIGAGHLFRLSG YPRDDQNREYLVVGAEYRVVQELYETGSGGAGSQFESELDCIDASQSFRLL PQTPVPVVRGPQTAVVVGPKGEEIWTDQYGRVKVHFHWDRHDQSNENSS CWIRVSQAWAGKNWGSMQIPRIGQEVIVSFLEGDPDRPIITGRVYNAEQTV PYELPANATQSGMKSRSSKGGTPANFNEIRMEDKKGAEQLYIHAERNQDN LVENDASLSVGHDRNKSIGHDELARIGNNRTRAVKLNDTLLVGGAKSDSVT GTYLIEAGAQIRLVCGKSVVEFNADGTINISGSAFNLYASGNGNIDTGGRLD LNSGGASEVDAKGKGVQGTIDGQVQAMFPPPAKGGSGSGSPSRLEEELRR RLTE

#### 1.2 SI-2: Microfluidic Device and Workflow

Description of the cryoWriter microfluidic platform, including the capillary system, magnetic trap, grid-writing workflow, robotic components, and control software.

The cryoWriter platform integrates microfluidic protein isolation with automated cryo-EM grid preparation. The system consists of a capillary-based microfluidic module with a magnetic trap for immobilizing superparamagnetic beads, a robotic positioning system for capillary and grid handling, and software for automated control of fluid handling and grid writing. Additional components provide UV-illumination for photocleavage and precise temperature control during grid preparation. For vitrification, strict temperature control of the grid and a ClimateJet [1] delivering an oxygen-free, temperature- and humidity-controlled gas stream are essential to ensure reproducible ice formation.

Figure 3 shows a photograph of the experimental setup and key components of the robotic system used for capillary manipulation and grid positioning.

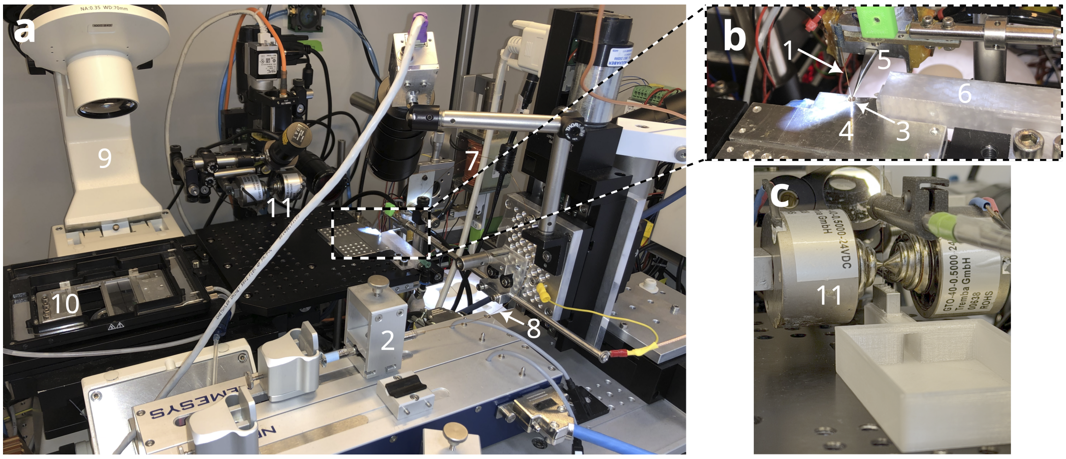

**Figure 3: Photograph and annotated overview of the cryoWriter platform: a)** Schematic overview of the cryoWriter modules. The microcapillary (1) connects the different functional units of the instrument. A high-precision pump (2) enables liquid handling with nanoliter precision. For grid preparation, a cryo-EM grid (3) is held in place on the dew-point stage (4) by tweezers (5). The ClimateJet [1] (6) controls the micro-climate around the grid and al-lows modulation of the air-water interface by injecting additives. After sample deposition, the grid is plunge-frozen (7) in liquid ethane (8). An inverted light microscope (9) and a cell-culturing compartment (10) enable experiments with live cells, such as single-cell lysis. The magnetic trap (11) allows immobilization of superparamagnetic beads for protein isolation and subsequent cryo-EM sample preparation. **b)** Close-up of the dew-point stage showing the capillary in the dispensing position above the grid, held by the temperature-controlled tweezer. The outlet of the ClimateJet directing the gas-stream to the grid is visible. **c)** Close-up of the magnetic trap module with inserted capillary and sample holder.

Figure 4 presents screenshots of the control software used to operate the microfluidic workflow, including capillary positioning, liquid handling, and automated grid deposition.

To prevent dilution of the eluted sample and to suppress Taylor dispersion within the capillary, the nanoliter-scale elution plug is separated from the system liquid by a small air bubble. Figure 5 illustrates a control experiment demonstrating the importance of introducing this air bubble prior to the isolation experiment. Pulling the air bubble across the bead bed after immobilization of the protein can damage sensitive protein assemblies, as shown here for apoferritin.

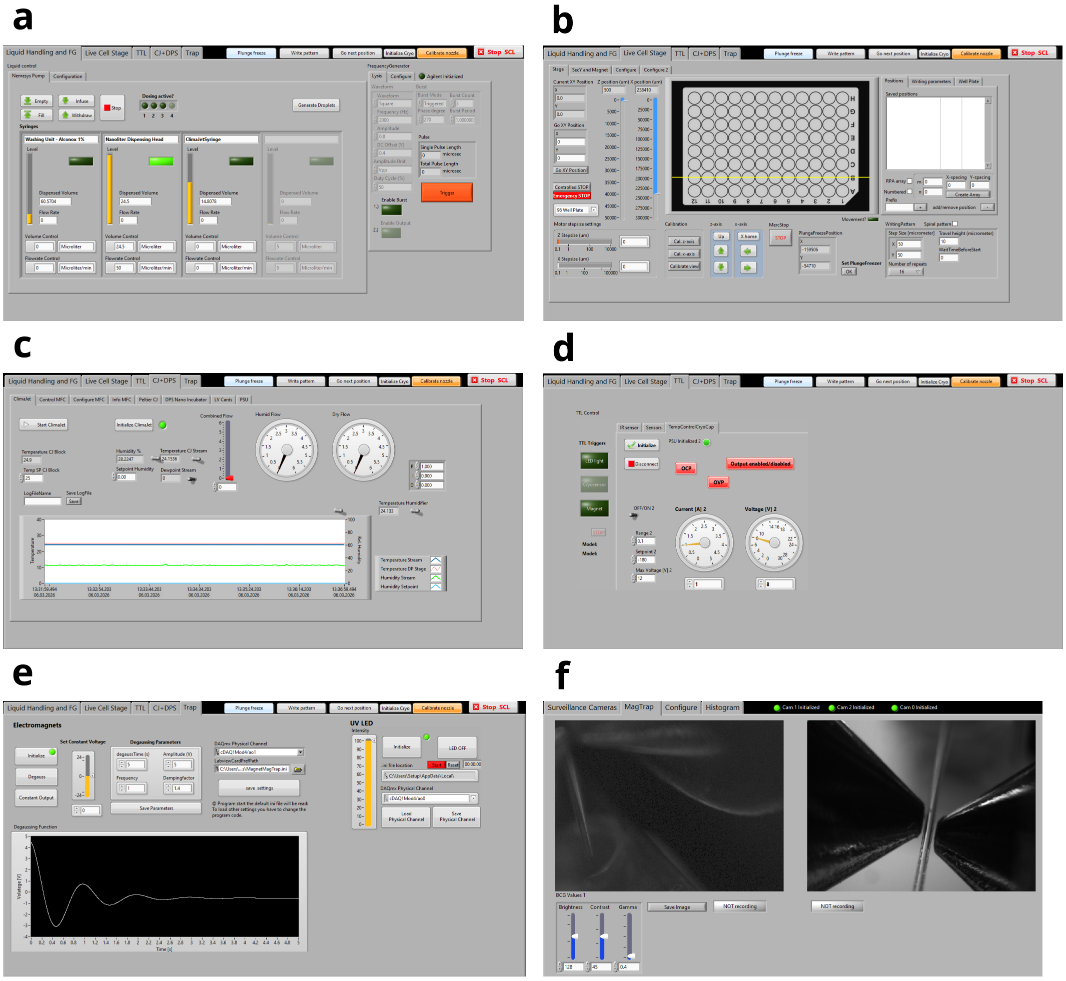

**Figure 4: CryoWriter control software implemented in the openBEB framework:** The cryoWriter platform is operated using the openBEB software environment [2], which was previously developed in our group. openBEB macros enable automation of cryoWriter workflows, including sample uptake, capillary positioning, grid deposition, and plunge-freezing. **a)** Liquid control panel for nanoliter-precise control of liquid movement within the capillary. **b)** Stage control panel for positioning the capillary, the translation stages, and the magnetic trap. **c)** ClimateJet panel for controlling the micro-climate around the grid, including the ClimateJet gas stream and the dew-point stage temperature. **d)** Control panel used to regulate the temperature of the liquid ethane pot. **e)** Magnetic trap control panel for adjusting the magnetic field strength and performing degaussing to remove residual magnetization of the magnetic trap. **f)** Live camera view used to monitor the capillary, dew-point stage, and magnetic trap during operation.

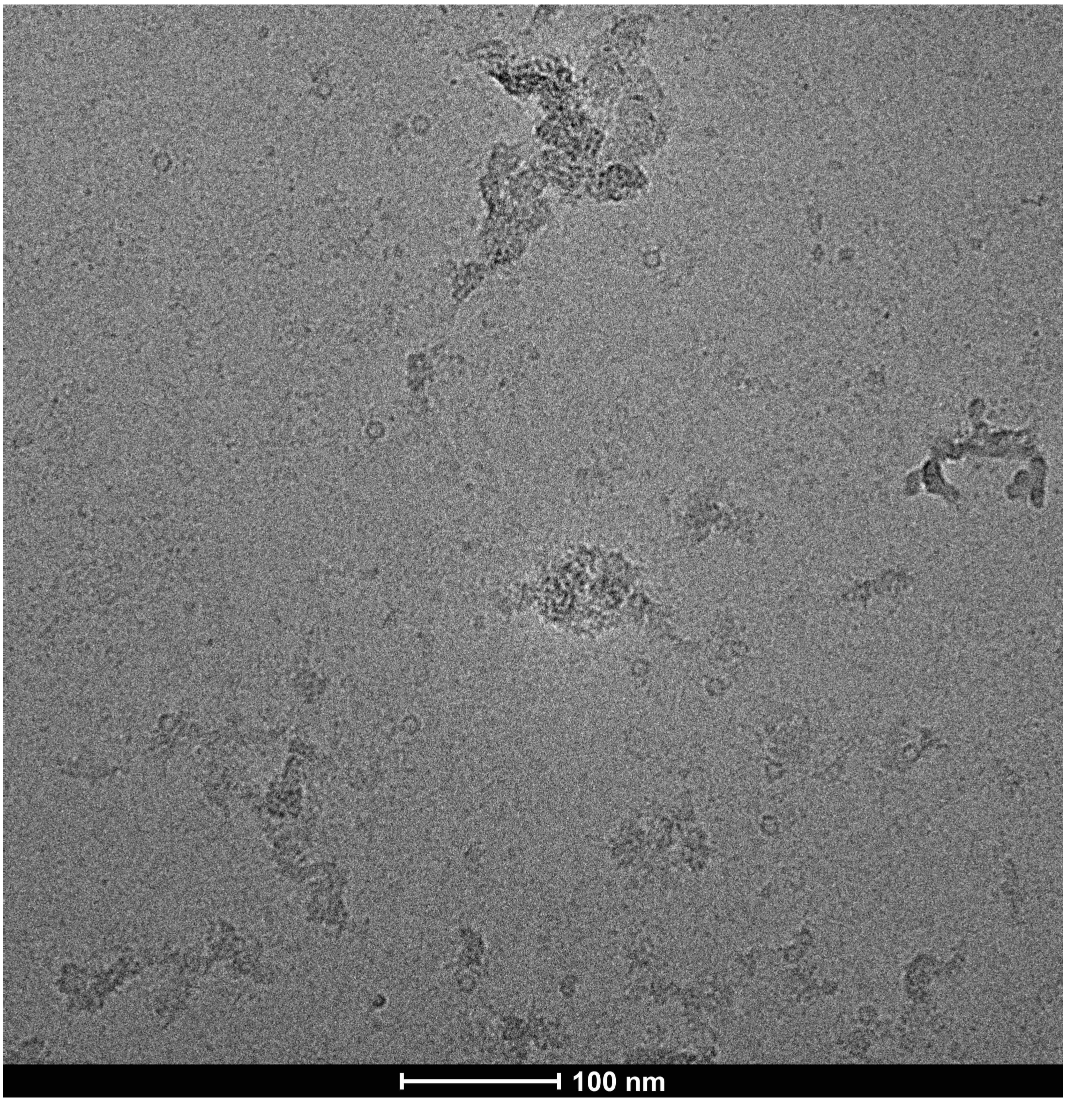

**Figure 5: Effect of pulling an air bubble across the bead plug:** Spy-ApoF isolated from cell lysate using a modified protocol in which an air bubble was pulled across the bead plug to enclose the beads in a small buffer volume prior to photocleavage. Only a small number of intact apoferritin particles are visible, while a large fraction of particles appears damaged or aggregated. This observation highlights the importance of introducing the air bubble before the isolation step rather than pulling it across the immobilized beads prior to photocleavage. Image recorded on a Talos microscope operated at 200 kV.

#### 1.3 SI-3: Tag Benchmarking

To evaluate the general applicability of the microfluidic isolation workflow, we compared several commonly used tagging strategies for affinity-based protein capture, including the $His_6$-tag, ALFA-tag, and the covalent SpyTag3/SpyCatcher3 system. These tagging strategies were assessed with respect to capture efficiency, compatibility with photocleavable linker chemistry, and suitability for downstream cryo-EM analysis. The following sections summarize the experimental observations for the different tagging strategies.

As discussed in the main manuscript, FLAG-tag capture was also explored during method development but showed reduced binding efficiency after crosslinker functionalization of the antibody and was therefore not pursued further.

##### 1.3.1 SI-3A: Microfluidic Isolation of His-tagged Proteins

His-tagging is one of the most widely used strategies for protein purification. In conventional workflows, His-tagged proteins are typically isolated using immobilized metal affinity chromatography (IMAC) followed by competitive elution and often an additional purification step such as size-exclusion chromatography.

To evaluate the suitability of His-tag capture for microfluidic batch isolation, we tested several capture strategies and constructs:

- His-tag-binding antibodies and derived Fab fragments.
- Photocleavable low-molecular-weight NTA derivatives. Two variants were tested, both showing similar limitations. Here we present the derivative that can be assembled via click chemistry from commercially available components.
- Aptamer-based capture reagents (the binding affinities obtained in our hands were too low for efficient isolation).

Across these approaches, three main limitations were observed.

- **Specificity:** Significant co-isolation of contaminant proteins was observed. Strategies commonly used in chromatographic purification, such as the addition of low concentrations of imidazole or detergents (e.g., Tween), reduced non-specific binding but also substantially decreased the yield.
- **Capture efficiency:** Although His-tag-binding antibodies exhibited good equilibrium binding affinities, the overall capture yields remained low under microfluidic batch conditions.
- **Interference with cryo-EM analysis:** Tris-NTA derivatives improved capture efficiency of $His_6$-tagged proteins; however, the release of low-molecular-weight components during photocleavage increased background signal and interfered with cryo-EM imaging (see Figure 6). To mitigate these effects, we reduced the density of photocleavable NTA crosslinkers on the bead surface by diluting them with standard biotin during the functionalization step. This modification improved the quality of the vitrified ice and reduced background signal in the micrographs, but at the cost of reduced particle yield.

Taken together, these observations indicate that while His-tags are highly effective in chromatographic purification workflows, they are less well suited for the microfluidic batch isolation strategy used here without further optimization of capture chemistry and surface functionalization.

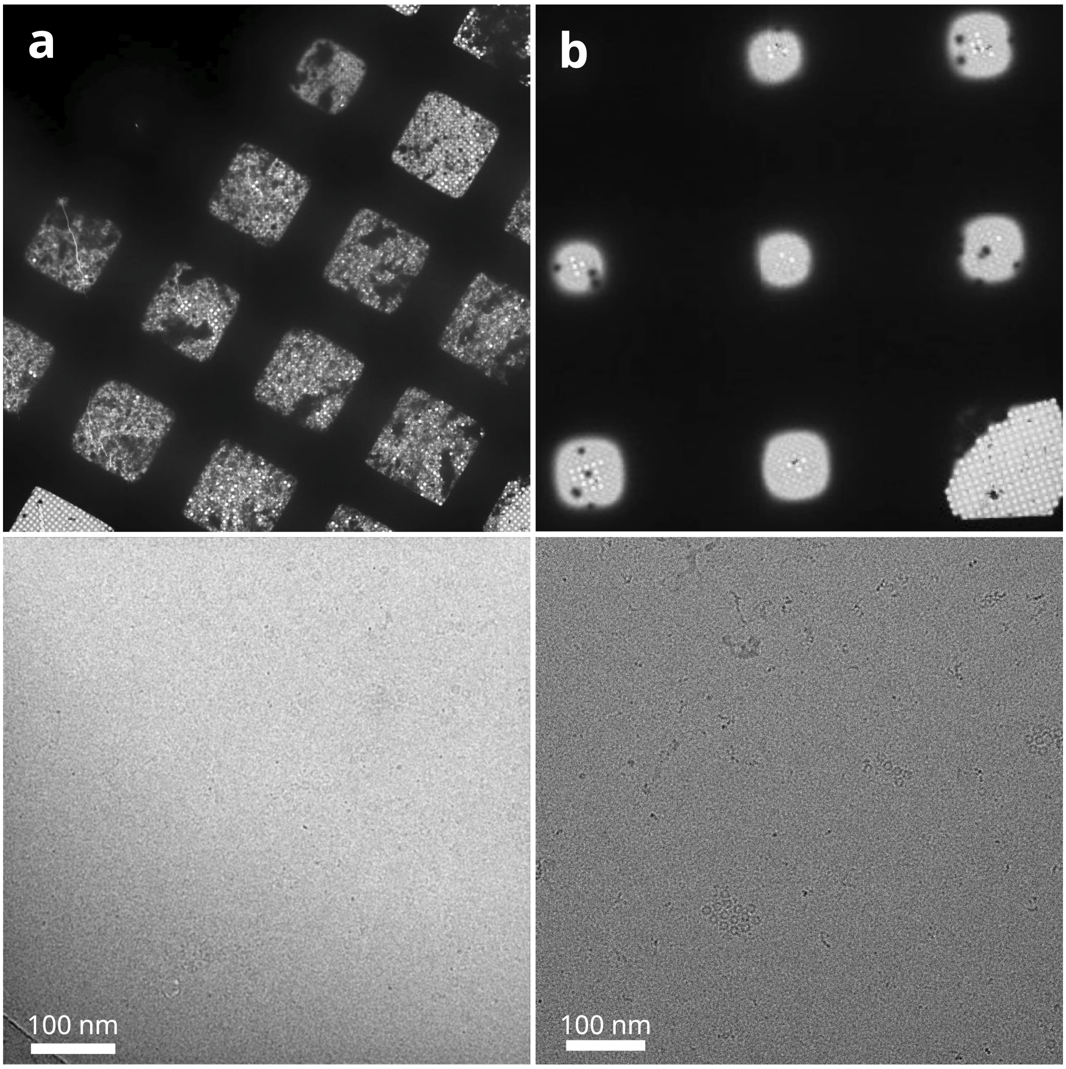

**Figure 6: Overview and close-up image of His-ALFA-ApoF isolation with the cryoWriter using the Tris-NTA-amine-crosslinker: a)** Here, a ratio of 1:10 (crosslinker:biotin) was used for functionalizing the beads in order to avoid an excess of bound crosslinker on the beads. Using this much crosslinker had detrimental effects on the ice quality, as can be seen from the dark areas on the atlas, which are caused by precipitated protein that was bound unspecifically to the NTA-crosslinker. Large amounts of free unbound NTA-crosslinker likely also had negative effects on the ice quality. The close-up image had to be low-pass filtered to 20 Å in order to better visualize the proteins. This was necessary due to the very low contrast which is likely caused by the presence of unbound NTA-crosslinker. While the amount of isolated protein was quite good, the subpar ice quality (blurry, low-contrast and rapid melting) prevented high-resolution data collection. **b)** Here, a ratio of 1:12 (crosslinker:biotin) was used for functionalizing the beads. Using less crosslinker led to better ice, but also fewer proteins. In addition, many holes did not have ice in them due to the low amount of protein. However, a balance between the amount of protein on the grid and acceptable ice quality could not be found.

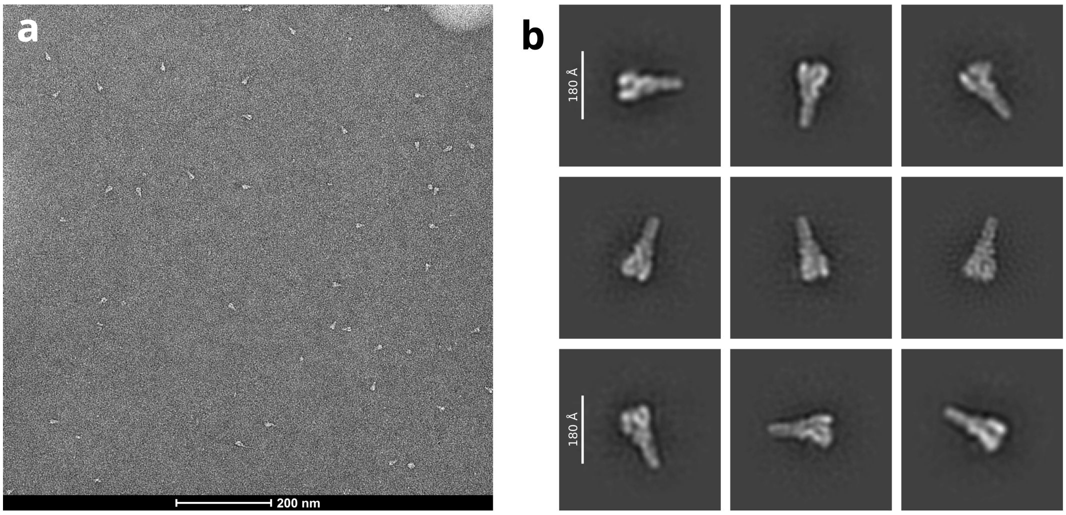

**Figure 7: Negative-stain characterization of VgrG1-His isolated using photocleavable Tris-NTA-amine crosslinkers: a)** Representative negative-stain micrograph of VgrG1-His following isolation with Tris-NTA-amine-functionalized beads. **b)** Representative 2D-class averages obtained from the negative-stain dataset. While VgrG1-His could be successfully captured and visualized by negative-stain EM, samples isolated using the Tris-NTA-amine crosslinker showed substantial aggregation under cryo-EM grid preparation conditions and did not yield datasets suitable for high-resolution reconstruction. In contrast, isolation of VgrG1 using the SpyTag3/SpyCatcher3 system produced well-dispersed particles and enabled high-resolution structure determination (see main text).

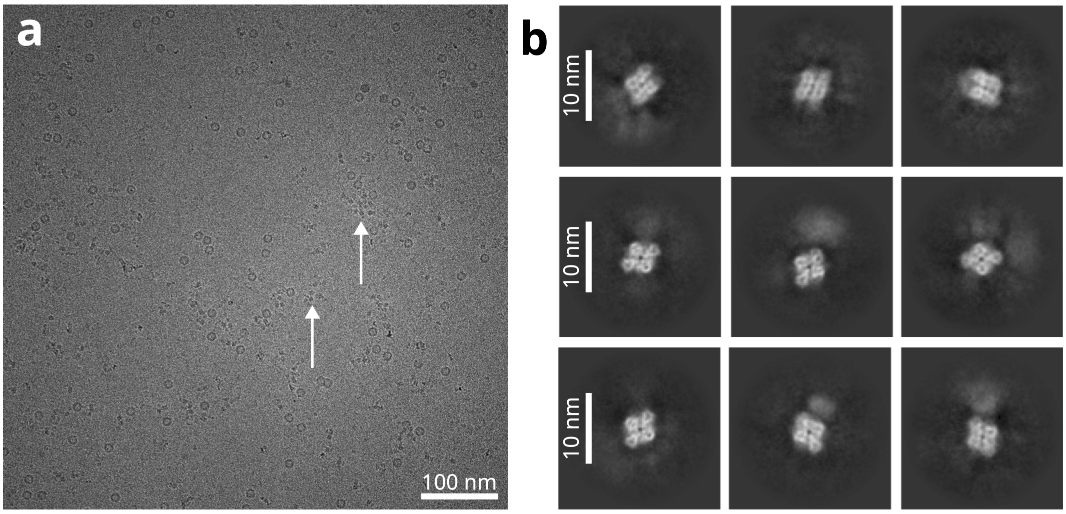

**Figure 8: Microfluidic isolation of Mat2a-His using anti-His Fab fragments: a)** Representative cryo-EM micrograph of Mat2a isolated from clarified cell lysate using anti-His Fab capture reagents. Mat2a particles are indicated by white arrows. Apoferritin (ApoF) was added to the sample to improve ice stability during grid preparation. The Mat2a-containing lysate was provided by collaborators at Roche. **b)** Representative 2D class averages of the isolated Mat2a particles. Weak peripheral density surrounding the protein likely corresponds to the bound Fab fragment. The dataset exhibited strong preferred particle orientation, resulting predominantly in top views and preventing reliable 3D reconstruction. The overall particle yield was lower than observed for other tagging strategies, likely reflecting the weaker binding characteristics of the anti-His Fab reagents under the microfluidic isolation conditions.

##### 1.3.2 SI-3B: Additional Microfluidic Isolation of SpyTag3- and ALFA-tagged Proteins

Constructs of $\beta$-galactosidase and VgrG1 carrying either SpyTag3 or the ALFA-tag were isolated using the cryoWriter. The resulting grids were screened by cryo-EM to assess their suitability for high-resolution data collection. Based on particle distribution and preliminary data quality, VgrG1-Spy and $\beta$-galactosidase-ALFA were selected for further data collection and reconstruction.

Representative micrographs of the two alternative constructs that were not pursued further are shown in Figure 9. Binding assays and additional biochemical characterizations of the constructs are provided in subsection 1.4.

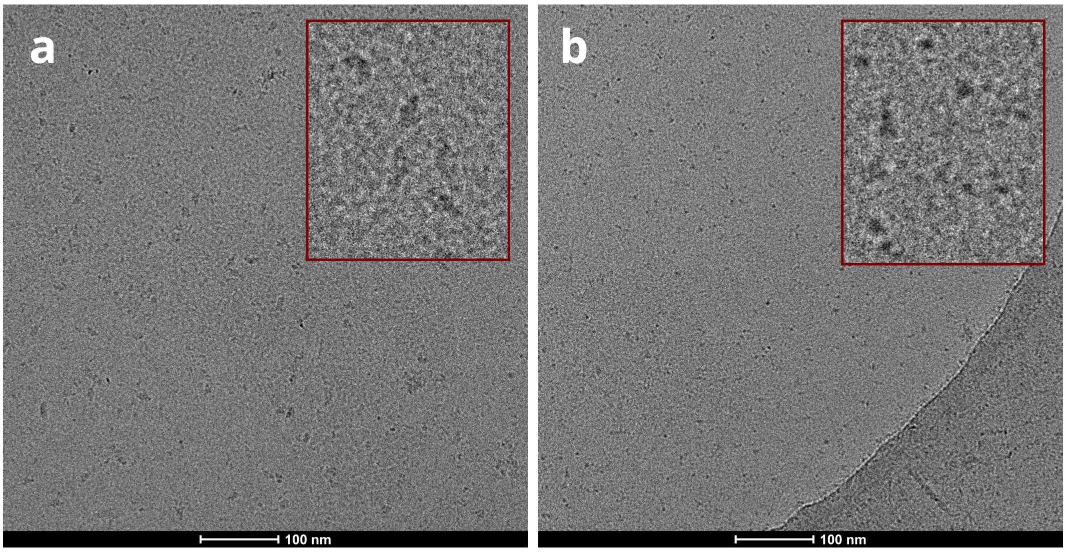

**Figure 9: Screening of additional isolated constructs: a)** Representative cryo-EM micrograph of $\beta$-galactosidase-SpyTag3 following microfluidic isolation. **b)** Representative cryo-EM micrograph of VgrG1-ALFA following microfluidic isolation. Both constructs showed good particle yield on the grids during screening. Images were recorded on a Talos microscope operated at 200 kV.

##### 1.3.3 SI-3C: Compatibility with GFP-tag capture

To explore whether commonly used fluorescent protein tags are compatible with the microfluidic isolation workflow, we tested GFP-tagged constructs using GFP-specific nanobodies as capture reagents. GFP tags are widely used for protein localization studies in living cells and therefore represent an attractive strategy for linking cellular imaging with structural analysis (see Discussion in the main manuscript).

GFP-tagged proteins could be successfully captured and isolated using GFP nanobody-functionalized beads within the microfluidic workflow. Negative-stain and cryo-EM screening confirmed that sufficient protein quantities could be recovered for imaging. Figure 10 shows the result of an experiment using GFP-nanobodies to capture and isolate GFP-tagged cold-shock DEAD-box protein A (CsdA) from cell lysate. Due to the inherent flexibility of the construct, only a low-resolution structure could be obtained, but the three main domains of the construct can be assigned by comparison with the AlphaFold-predicted structure. These results serve as a proof-of-concept for the compatibility of GFP tags with the microfluidic isolation workflow and it should be possible to obtain high-resolution structures from more suitable constructs in the future.

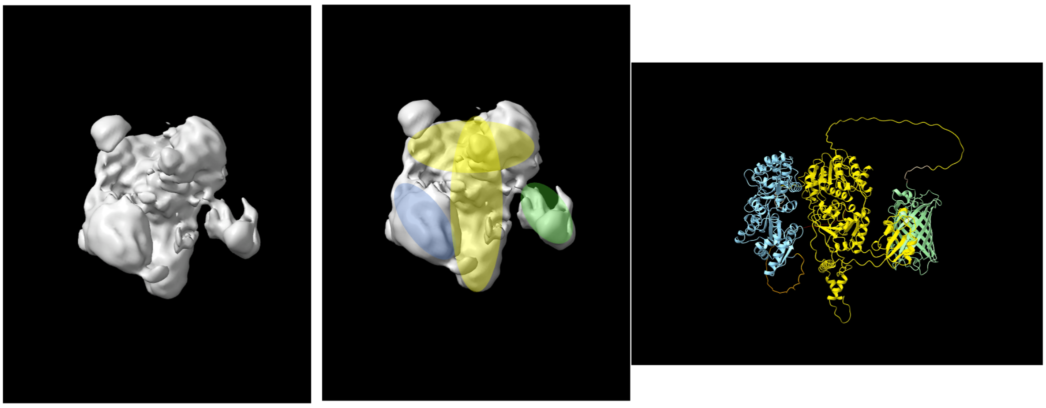

**Figure 10: Microfluidic isolation of GFP-tagged cold-shock DEAD-box protein A:** CsdA was isolated from cell lysate using GFP-nanobodies. This construct is made up of three main parts; CsdA (yellow), MBP (blue) and GFP (green). The isolation worked well, but the flexibility of the construct made 3D-reconstruction challenging, thus only a low-resolution structure could be obtained. Nevertheless, comparing the obtained volume to the AlphaFold-predicted structure (right panel) lets us assign the three domains to the volume.

In this work we focus primarily on proteins carrying short peptide tags. However, the experiments shown here demonstrate that GFP-tagged proteins can also be robustly isolated using the microfluidic workflow. Importantly, one of the tested constructs (Figure 11) represents a membrane protein, indicating that the isolation strategy is compatible with membrane-protein samples.

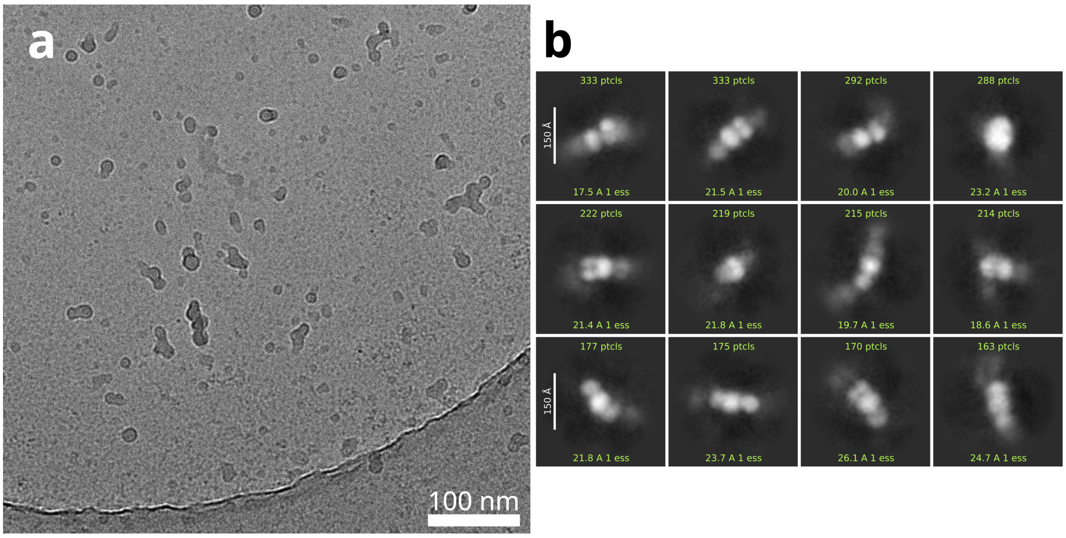

**Figure 11: Microfluidic isolation of a GFP-tagged ABC transporter using anti-GFP nanobodies: a)** Representative cryo-EM micrograph of the GFP-tagged ABC transporter following microfluidic isolation. The clarified lysate containing the target protein was provided by collaborators at Roche. **b)** Representative 2D class averages of the ABC transporter (scale bar: 15 nm). Only a small number of intact particles were observed on the grids, while the majority of complexes appeared partially dissociated during cryo-EM grid preparation. Consequently, the dataset was not suitable for 3D reconstruction. Nevertheless, the number of isolated particles indicates that the GFP-based capture strategy can provide sufficient material for cryo-EM analysis. The observed instability likely reflects the properties of this particular construct rather than a general limitation of the microfluidic isolation workflow.

#### 1.4 SI-4: Additional Tag Analysis and Control Experiments

Supporting biochemical characterization of the constructs and capture reagents, including pull-down assays analyzed by SDS-PAGE, bio-layer interferometry measurements, and additional validation experiments.

##### 1.4.1 Quality Control for Microfluidic Protein Isolation

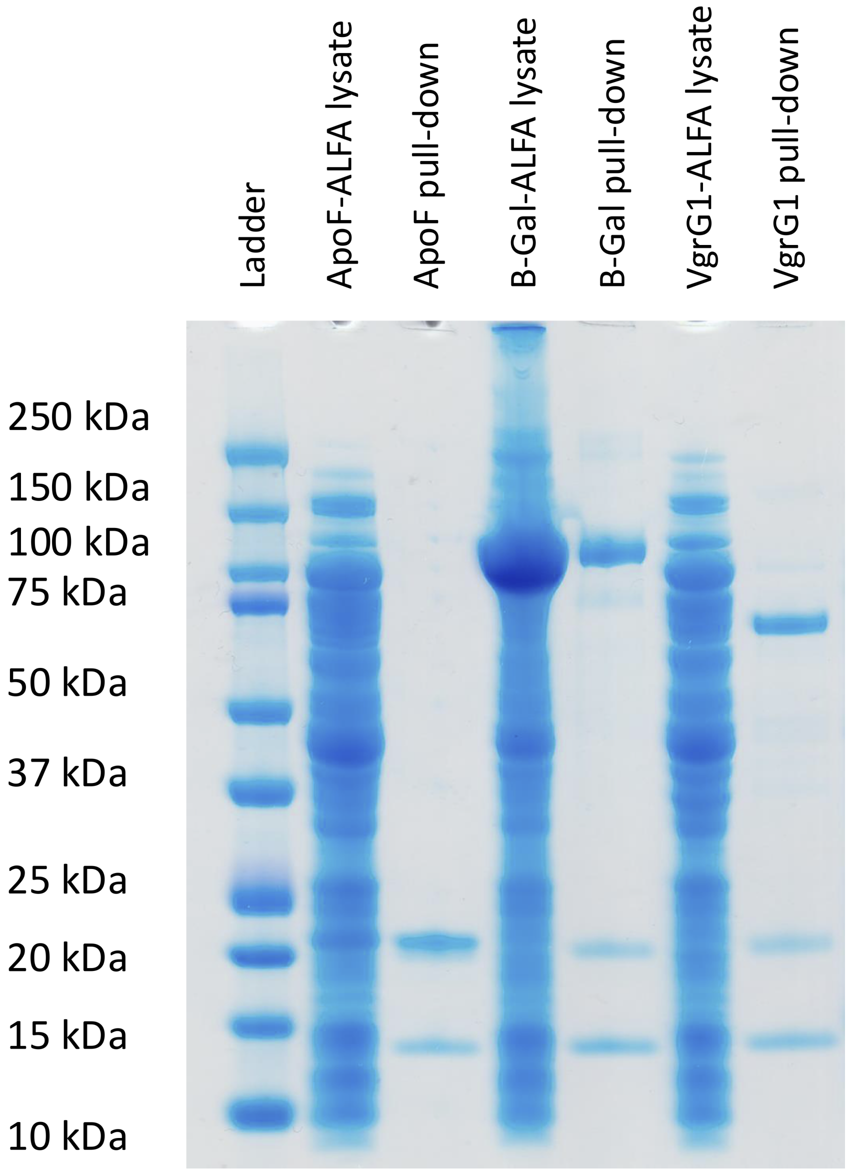

**Figure 12: Pull-down of ALFA-tagged ApoF, $\beta$-galactosidase and VgrG1:** ApoF and $\beta$-galactosidase can even be seen in the lane with the lysate due to their high level of expression. The band at around 13 kDa is due to the ALFA-nanobodies and the band at around 20 kDa is streptavidin from the streptavidin-coated magnetic beads used in the pull-down. ApoF, $\beta$-galactosidase and VgrG1 can be seen at the respective molecular weight for their monomers (23 kDa, 118 kDa and 74 kDa, respectively). There are basically no bands for unspecifically bound proteins, showing the specificity of the ALFA-nanobody, which makes it particularly well-suited for microfluidic protein isolation.

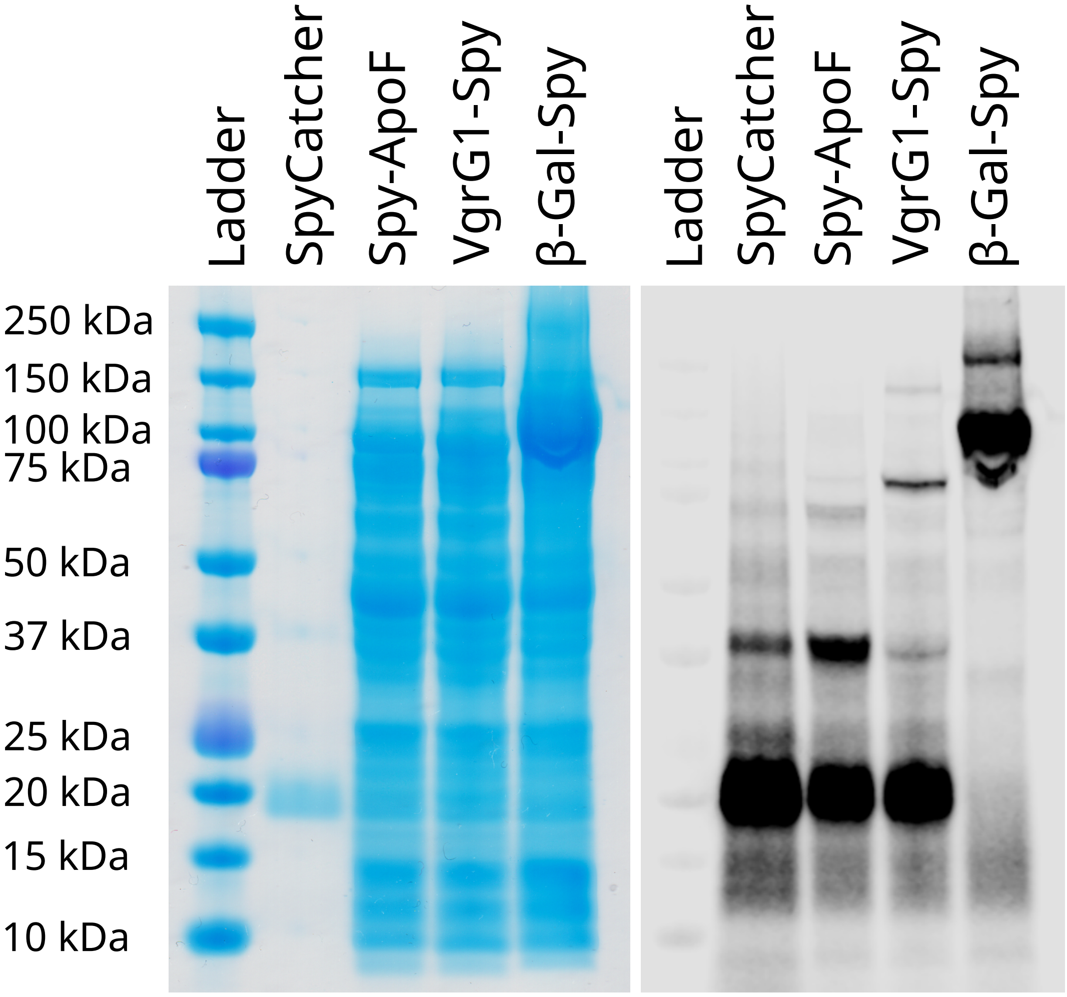

**Figure 13: SDS-PAGE gel of fluorescently labeled Spy-tagged proteins:** The lysates were incubated with fluorescently labelled SpyCatcher3, a gel was run and imaged with a fluorescence imager. This allows the fast detection of the Spy-tagged proteins without the need for purification. Bands can be seen for all Spy-tagged proteins at the molecular weight of their monomers plus the weight of the bound SpyCatcher3. I.e. the band for ApoF is at $\sim$35 kDa, for VgrG1 at $\sim$90 kDa and for $\beta$-galactosidase at $\sim$130 kDa. The band at $\sim$40 kDa in the SpyCatcher3-lane might be SpyCatcher3 that has dimerized via its cysteine residue and has been labelled at a lysine, rather than a cysteine. This band, along with the monomer SpyCatcher3-band, is absent in the $\beta$-galactosidase lane, but there is a second smaller band at $\sim$260 kDa, which corresponds to a SpyCatcher3 dimer with two bound $\beta$-galactosidase monomers. Similar, but much fainter bands like this can also be seen in the lanes of ApoF and VgrG1.

**Table 1: Concentration, $K_D$- and $R^2$-values for kinetic fits to ALFA-ApoF BLI data:** The fits for the measurements at 12.5 and 50 nM ALFA-ApoF were the best and the calculated $K_D$-values are likely the most accurate. The measurement device's sensitivity was a limiting factor for these measurements. Very low dissociation cannot be accurately fitted and thus an unrealistic $K_D$-value was calculated for the measurement using 200 nM ALFA-ApoF. However, these measurements, showing an affinity in the low nM-range, confirm the strong binding between the nanobodies and the tag, which is sufficient for our experiments.

| Concentration [nM] | Calculated $K_D$ [nM] | $R^2$-value |
|---|---|---|
| 12.5 | 4.4 | 0.9932 |
| 50 | 0.29 | 0.9997 |
| 200 | $< 0.001$ | 0.9768 |

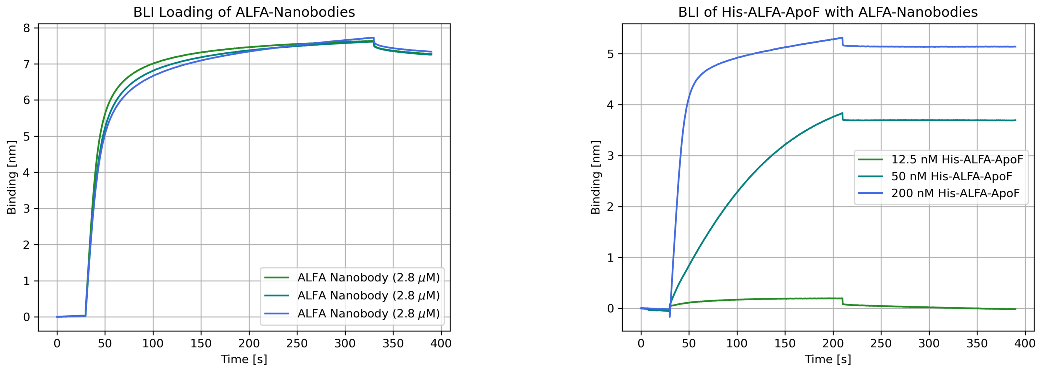

**Figure 14: Bio-layer interferometry measurements of ALFA-ApoF:** The dissociation constant $K_D$ of the ALFA-nanobodies towards ALFA-ApoF was measured. There was a clear signal for binding and there was only very little dissociation, leading to $K_D$-values in the low nM-range.

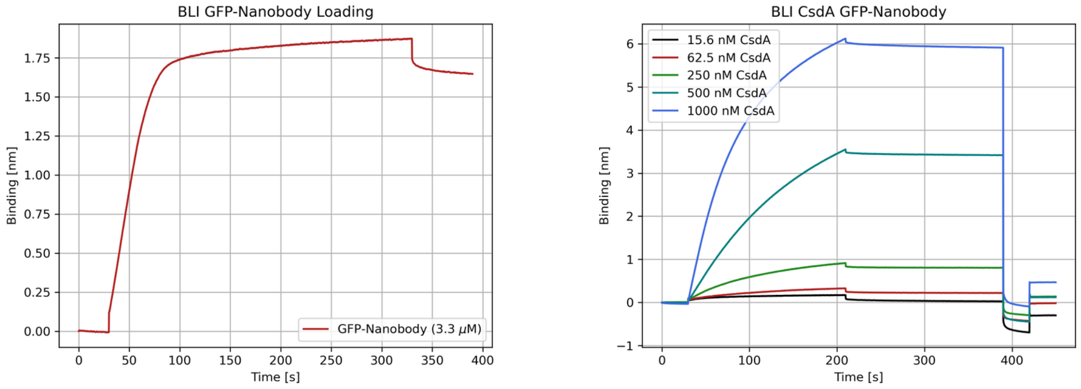

**Figure 15: Bio-layer interferometry measurements of CsdA:** Loading and measurement data of the GFP-nanobodies and CsdA construct. A $K_D$ of 2.8 nM was measured ($R^2 = 0.9998$). Importantly, there is very little dissociation, which makes the GFP-nanobody well-suited for protein isolation experiments.

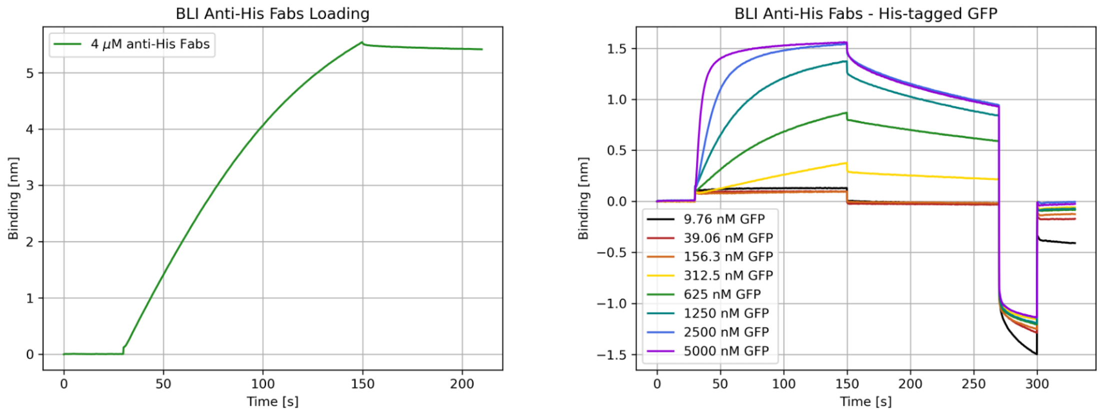

**Figure 16: Bio-layer interferometry measurements of His-GFP using anti-His-Fabs:** Loading and measurement data of the anti-His Fabs with His-tagged GFP. A $K_D$ of 135 nM was measured ($R^2$ = 0.9968). The off-rate is faster compared to the ALFA- and GFP-nanobodies, which is likely the reason for the poor performance of these Fabs.

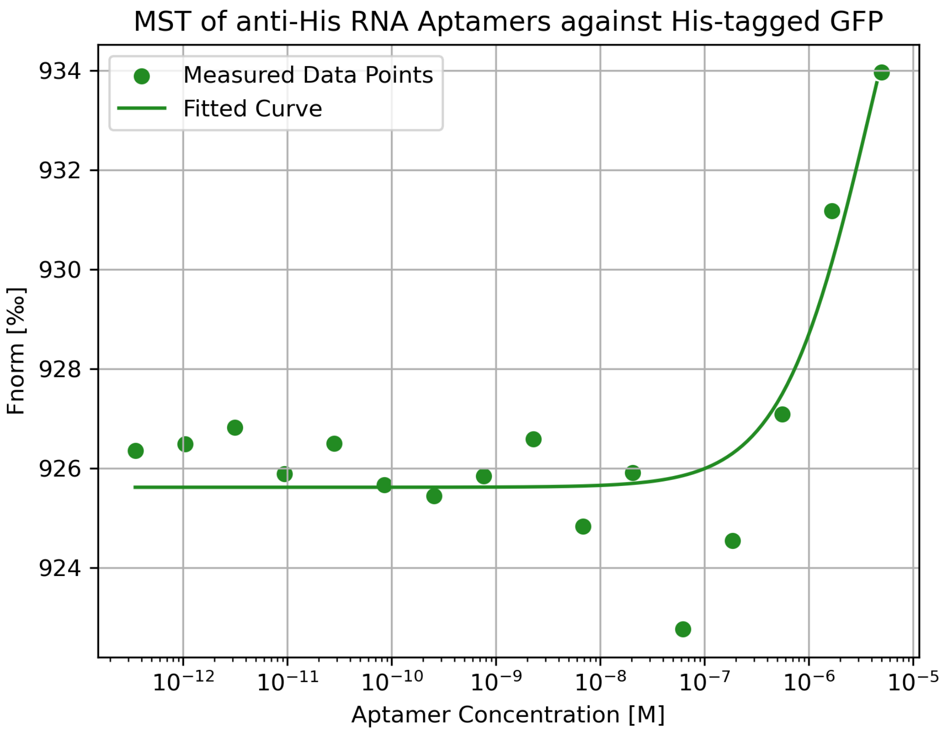

**Figure 17: Microscale Thermophoresis Measurement:** RNA-based aptamers binding His-tags [3] were measured to determine their binding strength. The measurement indicates weak binding ($K_D$ = 4.5±1 µM) which is insufficient for our needs.

#### 1.5 SI-5: Orientation Bias Analysis

Analysis of particle orientation distributions in cryo-EM datasets, including cFAR calculations, orientation plots, and structural analysis of affinity tags to rationalize differences in preferred orientation.

To analyze the influence of affinity tags on particle orientation in cryo-EM datasets, we compared the orientation distributions of particles isolated using the ALFA-tag and the SpyTag3/SpyCatcher3 system. Orientation distributions were evaluated using cFAR analysis and particle orientation plots.

We observed a pronounced difference in particle orientation distributions between ALFA-tagged and SpyTag3-tagged constructs (Figure 18); i.e. the ALFA-tag dataset exhibited a stronger preferred orientation compared to the SpyTag3 dataset. We attribute this difference to the distinct hydrophobicity patterns of the affinity tags (Figure 19), where the ALFA tag shows a more pronounced segregation of hydrophobic and hydrophilic regions that may promote interactions with the air-water interface.

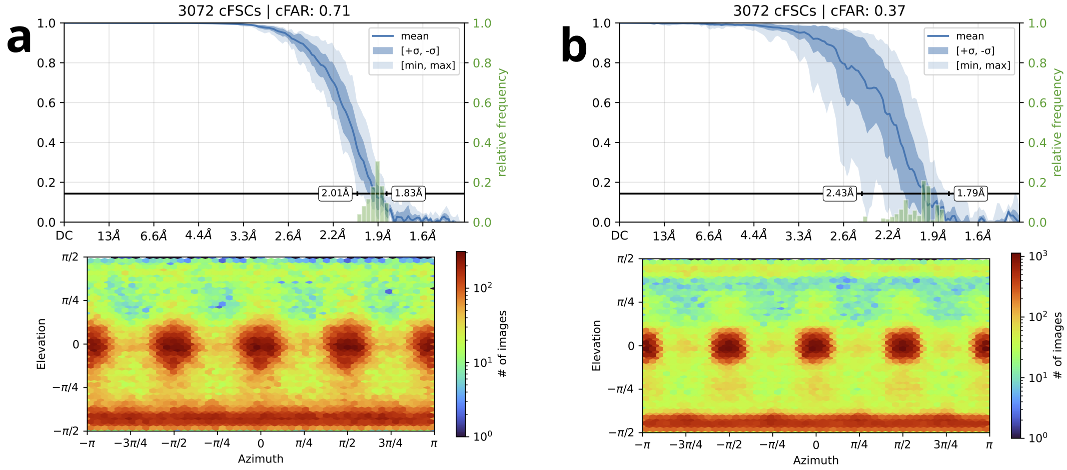

**Figure 18: cFAR and particle orientation distribution analysis: a)** Orientation distribution and cFAR analysis for Spy-ApoF isolated from cell lysate. The dataset shows a relatively uniform particle orientation distribution (cFAR = 0.71). **b)** Orientation distribution and cFAR analysis for ALFA-ApoF isolated from cell lysate. The dataset exhibits a stronger preferred orientation (cFAR = 0.37).

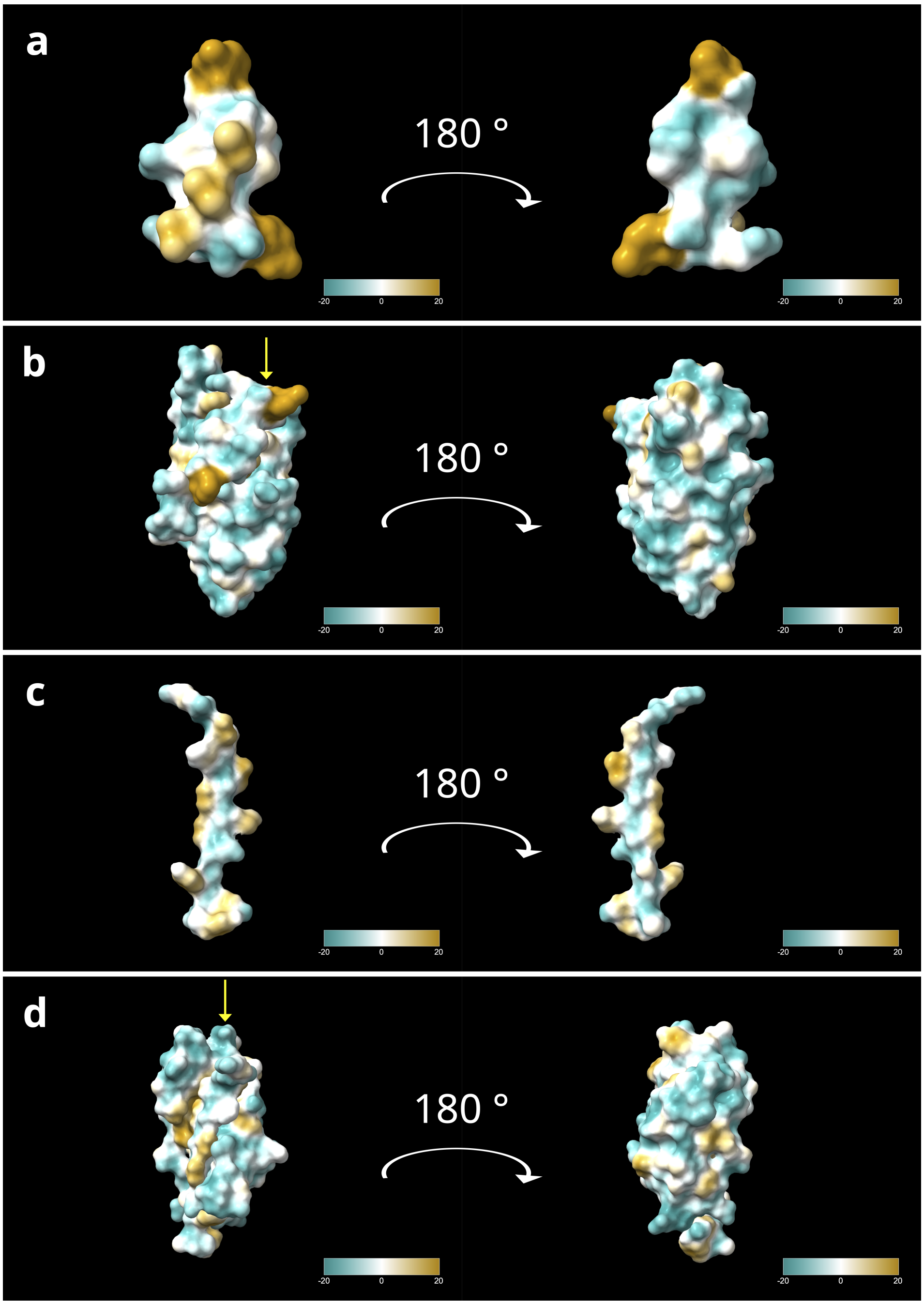

**Figure 19: Hydrophobicity analysis of affinity tags and capture proteins: a)** ALFA-tag (PDB: 6I2G) colored by hydrophobicity using the ChimeraX *mlp* command. The tag shows a segregation of hydrophobic and hydrophilic regions, with one side appearing more hydrophobic and the opposite side more hydrophilic. This asymmetry may contribute to the stronger preferred orientation observed for ApoF carrying an ALFA-tag compared with SpyTag3. **b)** ALFA-nanobody (theoretical pI 5.62) bound to the ALFA-tag (yellow arrow). The nanobody surface is predominantly hydrophilic and does not display large hydrophobic patches. **c)** SpyTag3 (PDB: 9OJ3) colored by hydrophobicity. Hydrophobic and hydrophilic regions are more evenly distributed across the tag surface, which may reduce interactions with the air-water interface. **d)** SpyCatcher3 (theoretical pI 4.23) bound to SpyTag3 (yellow arrow). Similar to the ALFA nanobody, the SpyCatcher3 surface is largely hydrophilic. Overall, the capture proteins appear predominantly hydrophilic, suggesting that the hydrophobic features of the affinity tags themselves, particularly the ALFA-tag, may play a major role in promoting preferred particle orientations during cryo-EM grid preparation.

#### 1.6 SI-6: Cryo-EM Data Processing and Model Validation

Detailed cryo-EM processing workflows, including particle statistics, FSC curves, reconstruction workflows, and model validation reports.

##### 1.6.1 ApoF Isolation from Cell Lysate

The processing workflows for the SpyTag3- and ALFA-tagged ApoF datasets are summarized in Figure 20 and Figure 21 and the corresponding PHENIX validation statistics for the final models are provided in Table 2 and Table 3, respectively.

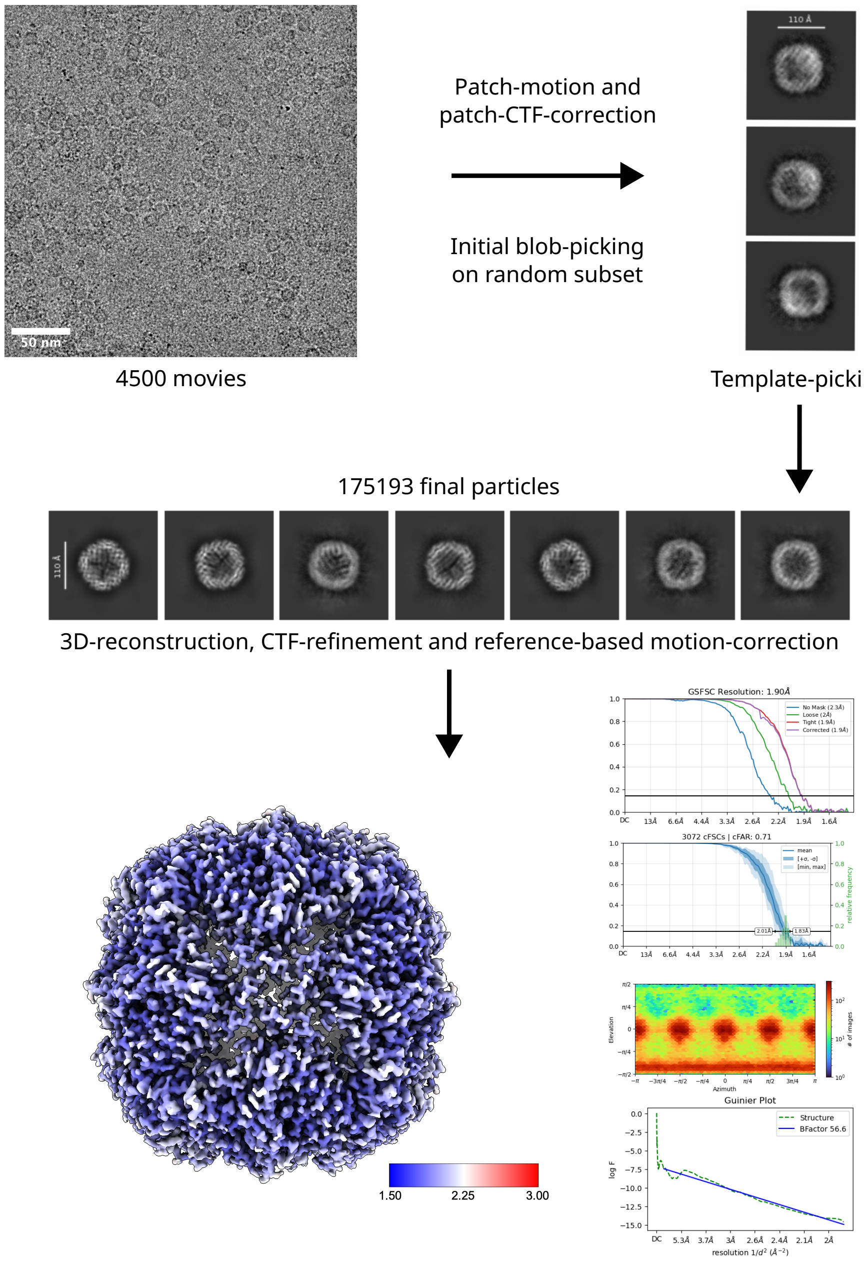

**Figure 20: Processing workflow of Spy-ApoF microfluidic protein isolation:** The image processing was carried out in CryoSPARC v4.7.1. The 4500 acquired movies were patch-motion- and patch-CTF-corrected. 4093 Micrographs with a CTF-fit of 7 Å or better were selected for further processing. Initial blob-picking (10-14 nm) was done on a random subset of 250 micrographs. 42 654 particles were picked and extracted with a box-size of 360 pixels, Fourier cropped to 180 pixels. These particles were 2D-classified into 50 classes and 3 high-quality classes were used for template-picking on the whole dataset. 629 947 particles were extracted with a box-size of 360 pixels, Fourier cropped to 180 pixels. Several rounds of 2D-classification were used to sort out low-quality particles, leading to a final 175 193 particles. These particles were extracted using a box-size of 360 pixels and an ab-inito model was calculated. This model was refined using the homogeneous refinement job with imposed octahedral symmetry as well as global and per-particle CTF-refinement, leading to a volume with a resolution of 2.04 Å. Lastly, the particles were subjected to reference based motion correction. The final homogeneous refinement job was performed with 152 336 particles and resulted in a volume with a resolution of 1.9 Å.

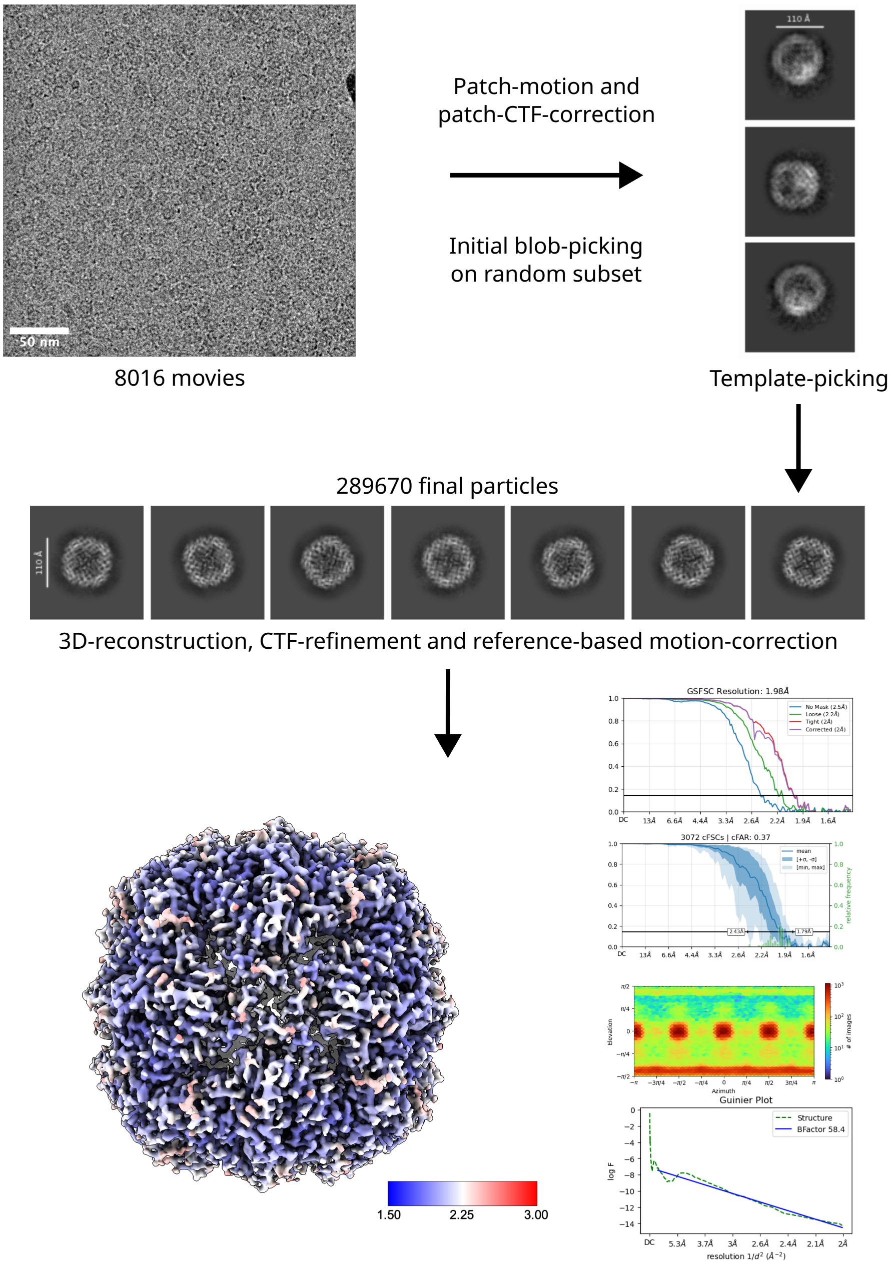

**Figure 21: Processing workflow of ALFA-ApoF microfluidic protein isolation:** The 8016 acquired movies were patch-motion- and patch-CTF-corrected. 7854 Micrographs with a CTF-fit of 7 Å or better were selected for further processing. Initial blob-picking (10-14 nm) was done on a random subset of 250 micrographs. 57 675 particles were picked and extracted with a box-size of 360 pixels, Fourier cropped to 180 pixels. These particles were 2D-classified into 50 classes and 3 high-quality classes were used for template-picking on the whole dataset. 1 430 697 particles were extracted with a box-size of 360 pixels, Fourier cropped to 180 pixels. Several rounds of 2D-classification were used to sort out low-quality particles, leading to a final 289 670 particles. These particles were extracted using a box-size of 360 pixels and an ab-inito model was calculated. This model was refined using the homogeneous refinement job with imposed octahedral symmetry as well as global and per-particle CTF-refinement, leading to a volume with a resolution of 2.04 Å. Lastly, the particles were subjected to reference based motion correction. The final homogeneous refinement job was performed with 289 367 particles and resulted in a volume with a resolution of 1.98 Å.

**Table 2:** PHENIX validation statistics for Spy-ApoF.

| | | |
|---|---|---|
| **Model** | | |
| Composition (#) | | |
| Chains | 24 | |
| Atoms | 32016 (Hydrogens: 0) | |
| Residues | Protein: 3864 Nucleotide: 0 | |
| Water | 0 | |
| Ligands | 0 | |
| Bonds (RMSD) | | |
| Length (Å) (# > 4$\sigma$) | 0.003 (0) | |
| Angles (°) (# > 4$\sigma$) | 0.475 (0) | |
| MolProbity score | 1.10 | |
| Clash score | 3.10 | |
| Ramachandran plot (%) | | |
| Outliers | 0.00 | |
| Allowed | 0.97 | |
| Favored | 99.03 | |
| "Rama-Z (Ramachandran plot Z-score RMSD)" | | |
| whole (N = 3816) | 5.26 (0.12) | |
| helix (N = 3312) | 3.76 (0.08) | |
| sheet (N = 0) | --- (---) | |
| loop (N = 504) | 1.03 (0.27) | |
| Rotamer outliers (%) | 0.77 | |
| C$\beta$ outliers (%) | NA | |
| Peptide plane (%) | | |
| Cis proline/general | 0.0/0.0 | |
| Twisted proline/general | 0.0/0.0 | |
| CaBLAM outliers (%) | 0.00 | |
| ADP (B-factors) | | |
| Iso/Aniso (#) | 32016/0 | |
| min/max/mean | | |
| Protein | 1.07/88.51/17.46 | |
| Nucleotide | --- | |
| Ligand | --- | |
| Water | --- | |
| Occupancy | | |
| Mean | 1.00 | |
| occ = 1 (%) | 100.00 | |
| 0 < occ < 1 (%) | 0.00 | |
| occ > 1 (%) | 0.00 | |
| **Data** | | |
| Box | | |
| Lengths (Å) | 130.67, 130.67, 130.67 | |
| Angles (°) | 90.00, 90.00, 90.00 | |
| Supplied Resolution (Å) | 1.9 | |
| Resolution Estimates (Å) | Masked | Unmasked |
| d FSC (half maps; 0.143) | --- | --- |
| d 99 (full/half1/half2) | 2.0/---/--- | 2.0/---/--- |
| d model | 2.0 | 2.0 |
| d FSC model (0/0.143/0.5) | 1.9/1.9/2.0 | 1.9/1.9/21.4 |
| Map min/max/mean | -0.22/0.40/0.00 | |
| **Model vs. Data** | | |
| CC (mask) | 0.88 | |
| CC (box) | 0.76 | |
| CC (peaks) | 0.76 | |
| CC (volume) | 0.85 | |
| Mean CC for ligands | --- | |

**Table 3:** PHENIX validation statistics for ALFA-ApoF.

| | | |
|---|---|---|
| **Model** | | |
| Composition (#) | | |
| Chains | 24 | |
| Atoms | 32016 (Hydrogens: 0) | |
| Residues | Protein: 3864 Nucleotide: 0 | |
| Water | 0 | |
| Ligands | 0 | |
| Bonds (RMSD) | | |
| Length (Å) (# > 4$\sigma$) | 0.006 (0) | |
| Angles (°) (# > 4$\sigma$) | 0.655 (0) | |
| MolProbity score | 1.49 | |
| Clash score | 4.48 | |
| Ramachandran plot (%) | | |
| Outliers | 0.00 | |
| Allowed | 0.73 | |
| Favored | 99.27 | |
| "Rama-Z (Ramachandran plot Z-score RMSD)" | | |
| whole (N = 3816) | 3.97 (0.12) | |
| helix (N = 3336) | 2.86 (0.08) | |
| sheet (N = 0) | --- (---) | |
| loop (N = 480) | 0.81 (0.27) | |
| Rotamer outliers (%) | 2.25 | |
| C$\beta$ outliers (%) | NA | |
| Peptide plane (%) | | |
| Cis proline/general | 0.0/0.0 | |
| Twisted proline/general | 0.0/0.0 | |
| CaBLAM outliers (%) | 0.00 | |
| ADP (B-factors) | | |
| Iso/Aniso (#) | 32016/0 | |
| min/max/mean | | |
| Protein | 6.06/53.31/19.50 | |
| Nucleotide | --- | |
| Ligand | --- | |
| Water | --- | |
| Occupancy | | |
| Mean | 1.00 | |
| occ = 1 (%) | 100.00 | |
| 0 < occ < 1 (%) | 0.00 | |
| occ > 1 (%) | 0.00 | |
| **Data** | | |
| Box | | |
| Lengths (Å) | 130.67, 129.21, 129.21 | |
| Angles (°) | 90.00, 90.00, 90.00 | |
| Supplied Resolution (Å) | 2.0 | |
| Resolution Estimates (Å) | Masked | Unmasked |
| d FSC (half maps; 0.143) | --- | --- |
| d 99 (full/half1/half2) | 2.1/---/--- | 2.1/---/--- |
| d model | 2.1 | 2.1 |
| d FSC model (0/0.143/0.5) | 1.9/1.9/18.0 | 1.9/2.0/18.0 |
| Map min/max/mean | -0.20/0.37/0.00 | |
| **Model vs. Data** | | |
| CC (mask) | 0.84 | |
| CC (box) | 0.71 | |
| CC (peaks) | 0.71 | |
| CC (volume) | 0.81 | |
| Mean CC for ligands | --- | |

##### 1.6.2 β-galactosidase Isolation from Cell Lysate

The 3D-reconstruction of β-galactosidase-ALFA colored by local resolution with fitted ALFA-nanobody is shown in Figure 22. Figure 23 shows the processing workflow for the β-galactosidase-ALFA dataset and the PHENIX validation statistics for the final model are provided in Table 4.

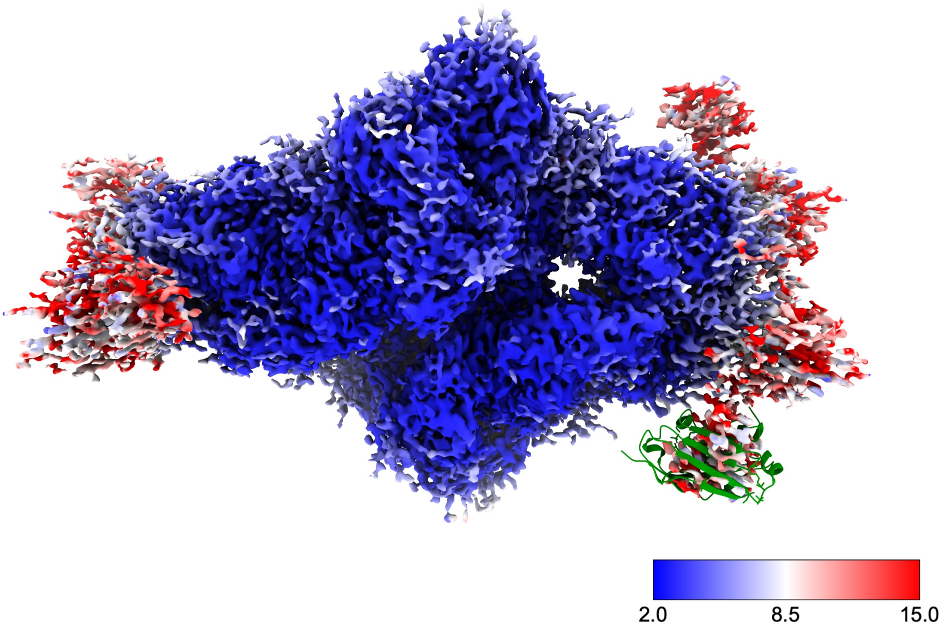

**Figure 22: 3D-reconstruction of β-galactosidase-ALFA colored by local resolution with fitted ALFA-nanobody:** The poorly resolved densities on the sides correspond to the bound ALFA-nanobody. There are several densities due to the imposed D2-symmetry and the flexibility of the linker sequence between β-galactosidase and the ALFA-tag. The structure of the ALFA-nanobody (PDB: 6I2G) has been manually placed in one of these densities in order to show that the size of the density matches the known structure. The bound nanobody did not negatively impact the reconstruction process of the target protein and high-resolution could be reached.

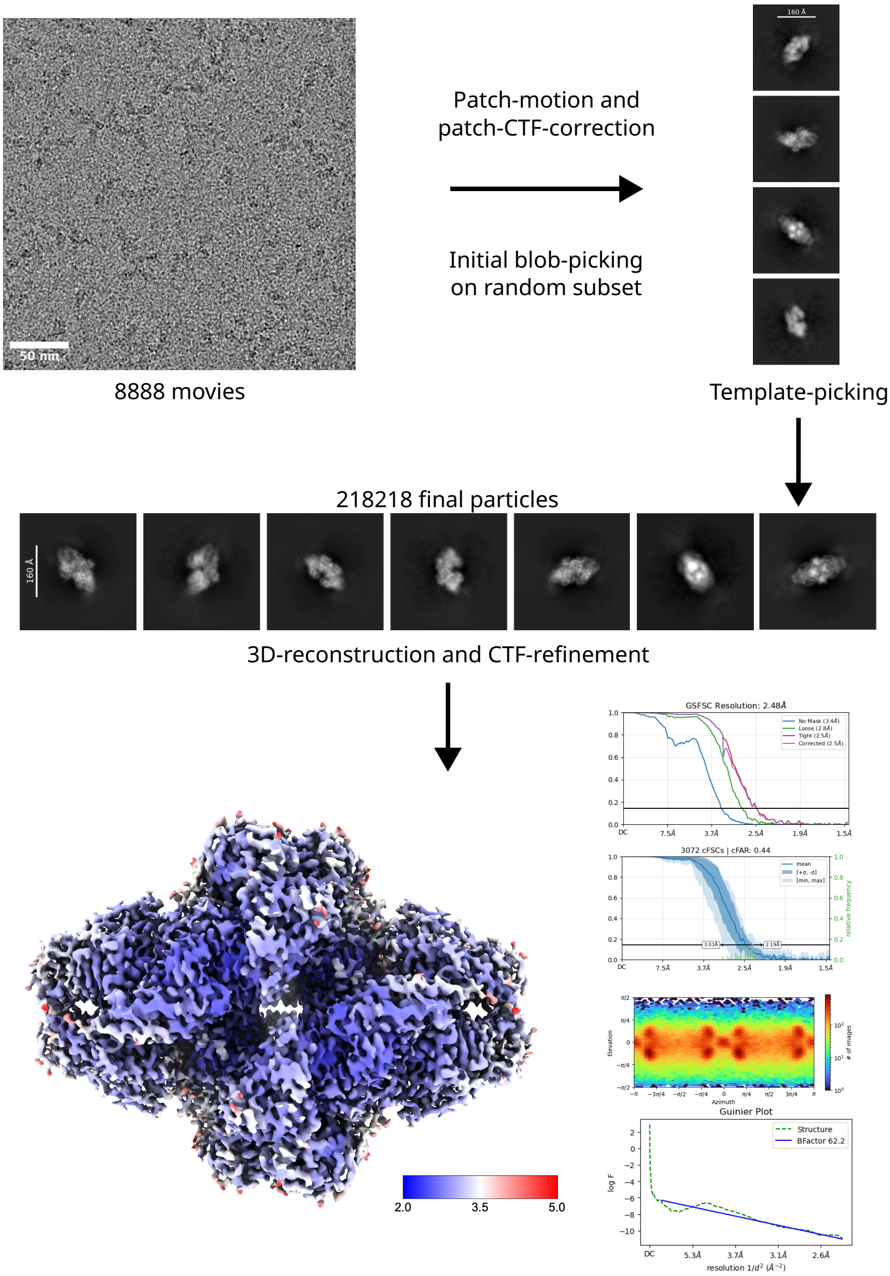

**Figure 23: Processing workflow of $\beta$-galactosidase-ALFA microfluidic protein isolation:** The 12 536 acquired movies were patch-motion- and patch-CTF-corrected. 8888 Micrographs with a CTF-fit of 7 Å or better and a relative ice-thickness below 1.3 were selected for further processing. Initial blob-picking (8-18 nm) was done on a random subset of 1000 micrographs. 66 255 particles were picked and extracted with a box-size of 512 pixels, Fourier cropped to 128 pixels. These particles were 2D-classified into 100 classes and 4 high-quality classes with different views were used for template-picking on the whole dataset. 814 849 particles were extracted with a box-size of 512 pixels, Fourier cropped to 128 pixels. Several rounds of 2D-classification were used to sort out low-quality particles, leading to a final 218 218 particles. These particles were extracted using a box-size of 512 pixels and an ab-inito model was calculated. This model was refined using the homogeneous refinement job with imposed D2-symmetry and global and per-particle CTF-refinement, leading to a volume with a resolution of 2.57 Å. This volume was used as input for a local refinement job, which resulted in a final volume with a resolution of 2.48 Å.

**Table 4:** PHENIX validation statistics for $\beta$-galactosidase-ALFA. Note that the published structure of *E.Coli* $\beta$-galactosidase (PDB:6drv) also has 8.1 % cis-Prolines.

| | | |
|---|---|---|
| **Model** | | |
| Composition (#) | | |
| Chains | 4 | |
| Atoms | 32872 (Hydrogens: 0) | |
| Residues | Protein: 4096 Nucleotide: 0 | |
| Water | 0 | |
| Ligands | 0 | |
| Bonds (RMSD) | | |
| Length (Å) (# > $4\sigma$) | 0.002 (0) | |
| Angles (°) (# > $4\sigma$) | 0.515 (0) | |
| MolProbity score | 1.53 | |
| Clash score | 3.96 | |
| Ramachandran plot (%) | | |
| Outliers | 0.00 | |
| Allowed | 2.57 | |
| Favored | 97.43 | |
| "Rama-Z (Ramachandran plot Z-score RMSD)" | | |
| whole (N = 4088) | 0.51 (0.13) | |
| helix (N = 552) | 0.36 (0.23) | |
| sheet (N = 1312) | 0.94 (0.15) | |
| loop (N = 2224) | 0.05 (0.13) | |
| Rotamer outliers (%) | 2.03 | |
| C$\beta$ outliers (%) | NA | |
| Peptide plane (%) | | |
| Cis proline/general | 8.1/0.3 | |
| Twisted proline/general | 0.0/0.0 | |
| CaBLAM outliers (%) | 2.50 | |
| ADP (B-factors) | | |
| Iso/Aniso (#) | 32872/0 | |
| min/max/mean | | |
| Protein | 9.59/160.38/66.57 | |
| Nucleotide | --- | |
| Ligand | --- | |
| Water | --- | |
| Occupancy | | |
| Mean | 1.00 | |
| occ = 1 (%) | 100.00 | |
| 0 < occ < 1 (%) | 0.00 | |
| occ > 1 (%) | 0.00 | |
| **Data** | | |
| Box | | |
| Lengths (Å) | 154.03, 194.91, 101.47 | |
| Angles (°) | 90.00, 90.00, 90.00 | |
| Supplied Resolution (Å) | 2.5 | |
| Resolution Estimates (Å) | Masked | Unmasked |
| d FSC (half maps; 0.143) | --- | --- |
| d 99 (full/half1/half2) | 2.8/---/--- | 2.7/---/--- |
| d model | 2.8 | 2.8 |
| d FSC model (0/0.143/0.5) | 2.4/2.5/2.7 | 2.5/2.5/2.9 |
| Map min/max/mean | -0.56/0.89/0.01 | |
| **Model vs. Data** | | |
| CC (mask) | 0.85 | |
| CC (box) | 0.68 | |
| CC (peaks) | 0.66 | |
| CC (volume) | 0.82 | |
| Mean CC for ligands | --- | |

##### 1.6.3 VgrG1 Isolation from Cell Lysate

The 3D-reconstruction of VgrG1-Spy colored by local resolution with fitted SpyCatcher3 is shown in Figure 24. The processing workflow for the VgrG1-Spy dataset is summarized in Figure 25 and PHENIX validation statistics for the final model are provided in Table 5.

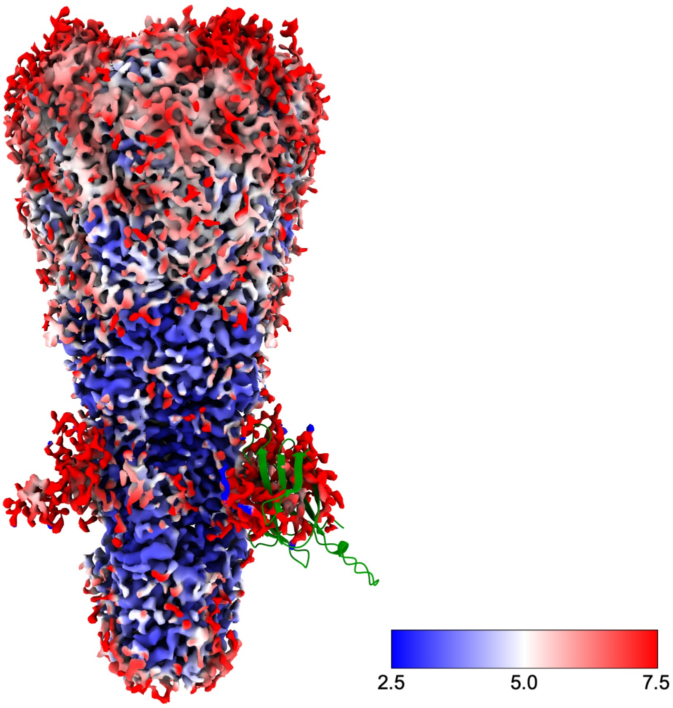

**Figure 24: 3D-reconstruction of VgrG1-Spy colored by local resolution with fitted SpyCatcher3:** The poorly resolved densities on the lower sides correspond to the bound SpyCatcher3. There are 3 densities due to the imposed C3-symmetry. SpyCatcher3 is not well resolved due to the flexibility of the linker sequence between VgrG1 and the SpyTag3. The AlphaFold3-predicted structure of SpyCatcher3-Cys has been manually placed in one of these densities in order to show that the size of the density matches the known structure. However, the resolution of the bound SpyCatcher3 was not good enough to resolve it. Nevertheless, the structure of VgrG1 could be refined to high-resolution without any issues caused by the bound SpyCatcher3, as shown in Figure 21.

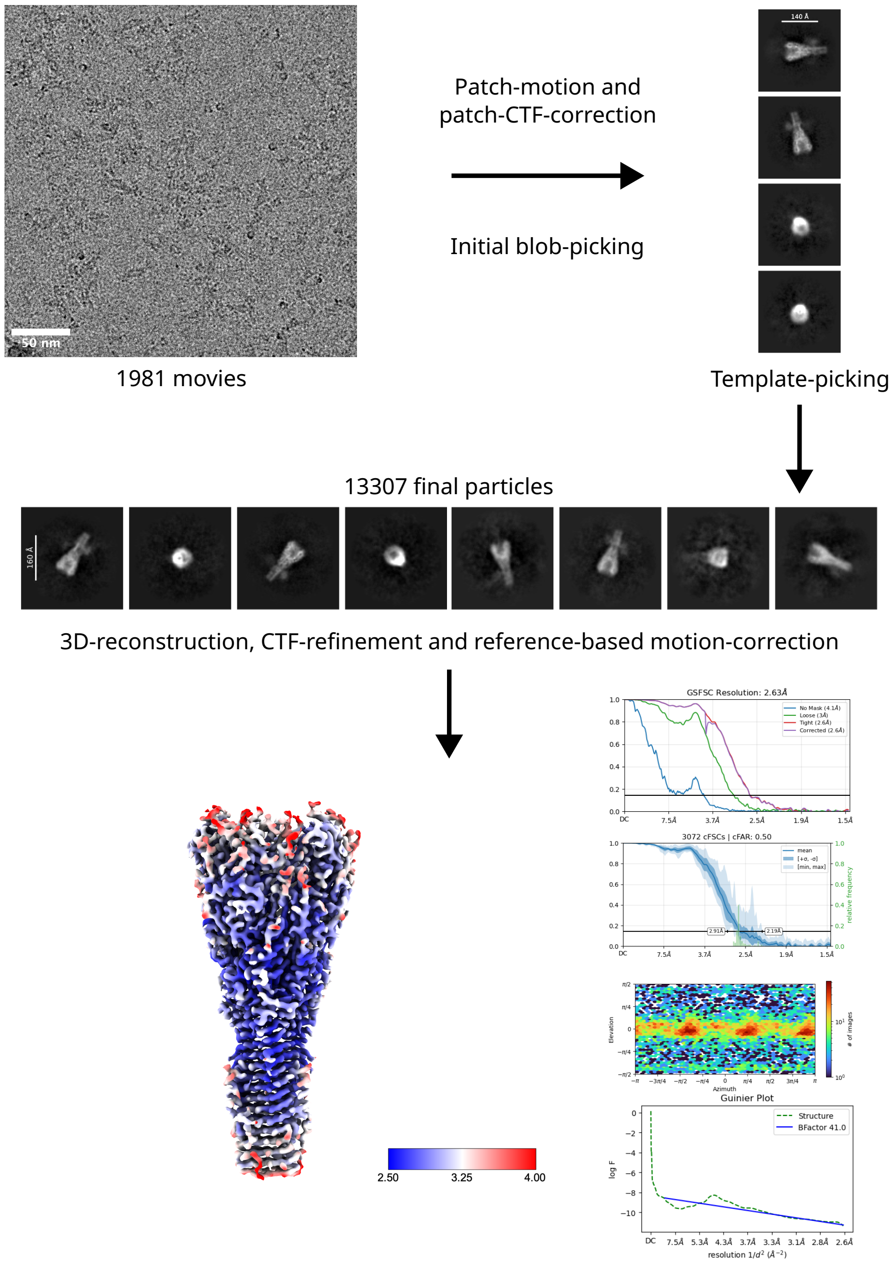

**Figure 25: Processing workflow of VgrG1-Spy microfluidic protein isolation:** The 3544 acquired movies were patch-motion- and patch-CTF-corrected. 1981 Micrographs with a CTF-fit of 7 Å or better were selected for further processing. Initial blob-picking (5-15 nm) was done on all micrographs. 175 428 particles were picked and extracted with a box-size of 448 pixels, Fourier cropped to 112 pixels. These particles were 2D-classified into 100 classes and 4 high-quality classes with a top- or side-view were used for template-picking on the whole dataset. 80 107 particles were extracted with a box-size of 448 pixels, Fourier cropped to 112 pixels. Several rounds of 2D-classification were used to sort out low-quality particles, leading to a final 13 307 particles. These particles were extracted using a box-size of 512 pixels and an ab-inito model was calculated. This model was refined using the homogeneous refinement job with imposed C3-symmetry, leading to a volume with a resolution of 4.06 Å. This volume was used as input for a non-uniform refinement job with global and per-particle CTF-refinement. Lastly, the particles were subjected to reference based motion correction. The final non-uniform refinement job was performed with 13 092 particles and resulted in a volume with a resolution of 2.63 Å.

**Table 5:** PHENIX validation statistics for VgrG1-Spy.

| | | |
|---|---|---|
| **Model** | | |
| Composition (#) | | |
| Chains | 3 | |
| Atoms | 15234 (Hydrogens: 0) | |
| Residues | Protein: 1926 Nucleotide: 0 | |
| Water | 0 | |
| Ligands | 0 | |
| Bonds (RMSD) | | |
| Length (Å) (# > $4\sigma$) | 0.003 (0) | |
| Angles (°) (# > $4\sigma$) | 0.521 (0) | |
| MolProbity score | 1.50 | |
| Clash score | 4.81 | |
| Ramachandran plot (%) | | |
| Outliers | 0.00 | |
| Allowed | 2.40 | |
| Favored | 97.60 | |
| "Rama-Z (Ramachandran plot Z-score RMSD)" | | |
| whole (N = 1920) | 0.23 (0.19) | |
| helix (N = 177) | 2.46 (0.40) | |
| sheet (N = 570) | -0.30 (0.22) | |
| loop (N = 1173) | 0.14 (0.18) | |
| Rotamer outliers (%) | 1.68 | |
| C$\beta$ outliers (%) | NA | |
| Peptide plane (%) | | |
| Cis proline/general | 5.9/0.2 | |
| Twisted proline/general | 0.0/0.0 | |
| CaBLAM outliers (%) | 1.57 | |
| ADP (B-factors) | | |
| Iso/Aniso (#) | 15234/0 | |
| min/max/mean | | |
| Protein | 26.81/187.71/73.67 | |
| Nucleotide | --- | |
| Ligand | --- | |
| Water | --- | |
| Occupancy | | |
| Mean | 1.00 | |
| occ = 1 (%) | 100.00 | |
| 0 < occ < 1 (%) | 0.00 | |
| occ > 1 (%) | 0.00 | |
| **Data** | | |
| Box | | |
| Lengths (Å) | 91.98, 91.25, 185.42 | |
| Angles (°) | 90.00, 90.00, 90.00 | |
| Supplied Resolution (Å) | 2.6 | |
| Resolution Estimates (Å) | Masked | Unmasked |
| d FSC (half maps; 0.143) | --- | --- |
| d 99 (full/half1/half2) | 2.9/---/--- | 2.8/---/--- |
| d model | 2.9 | 2.9 |
| d FSC model (0/0.143/0.5) | 2.5/2.6/2.8 | 2.6/2.6/3.1 |
| Map min/max/mean | -0.13/0.24/0.00 | |
| **Model vs. Data** | | |
| CC (mask) | 0.88 | |
| CC (box) | 0.69 | |
| CC (peaks) | 0.66 | |
| CC (volume) | 0.86 | |
| Mean CC for ligands | --- | |

##### 1.6.4 ApoF Isolation from IVT

The processing workflow for the ALFA-ApoF-IVT dataset is summarized in Figure 26. The model validation statistics for the final ALFA-ApoF-IVT model are provided in Table 6.

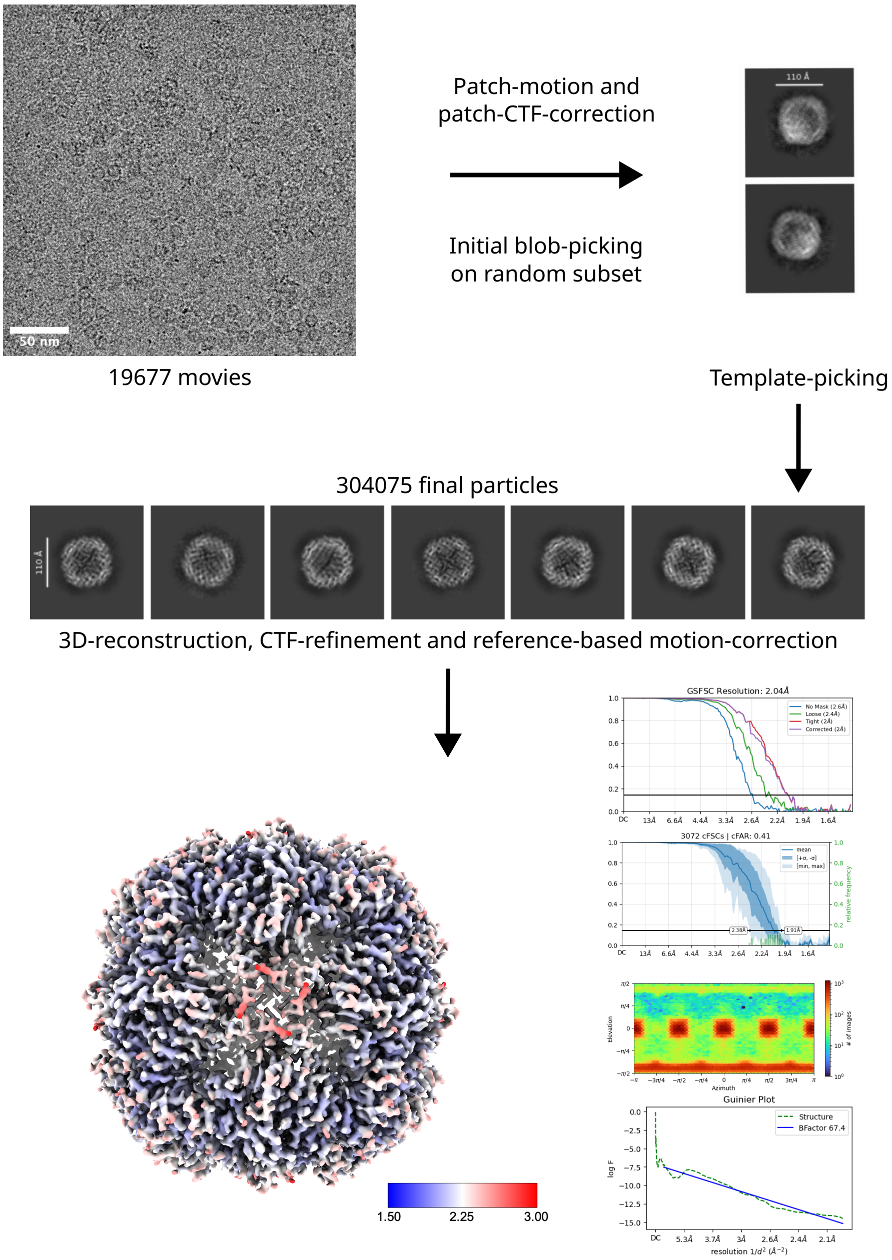

**Figure 26: Processing workflow of ALFA-ApoF-IVT microfluidic protein isolation:** The 19 677 acquired movies were patch-motion- and patch-CTF-corrected. 19 415 Micrographs with a CTF-fit of 7 Å or better were selected for further processing. Initial blob-picking (10-14 nm) was done on a random subset of 2000 micrographs. 350 480 particles were picked and extracted with a box-size of 360 pixels, Fourier cropped to 180 pixels. These particles were 2D-classified into 50 classes and classes containing ApoF-particles were classified again into 25 classes. 2 high-quality classes were used for template-picking on the whole dataset. 3 407 631 particles were extracted with a box-size of 360 pixels, Fourier cropped to 180 pixels. Several rounds of 2D-classification were used to sort out low-quality particles, leading to a final 305 365 particles. These particles were extracted using a box-size of 360 pixels and an ab-inito model was calculated. This model was refined using the homogeneous refinement job with imposed octahedral symmetry as well as global and per-particle CTF-refinement, leading to a volume with a resolution of 2.14 Å. Lastly, the particles were subjected to reference based motion correction. The final homogeneous refinement job was performed with 304 075 particles and resulted in a volume with a resolution of 2.04 Å.

**Table 6:** PHENIX validation statistics for ALFA-ApoF-IVT.

| | | |
|---|---|---|
| **Model** | | |
| Composition (#) | | |
| Chains | 24 | |
| Atoms | 32016 (Hydrogens: 0) | |
| Residues | Protein: 3864 Nucleotide: 0 | |
| Water | 0 | |
| Ligands | 0 | |
| Bonds (RMSD) | | |
| Length (Å) (# > 4$\sigma$) | 0.005 (0) | |
| Angles (°) (# > 4$\sigma$) | 0.873 (0) | |
| MolProbity score | 1.68 | |
| Clash score | 4.75 | |
| Ramachandran plot (%) | | |
| Outliers | 0.00 | |
| Allowed | 1.23 | |
| Favored | 98.77 | |
| "Rama-Z (Ramachandran plot Z-score RMSD)" | | |
| whole (N = 3816) | 5.20 (0.12) | |
| helix (N = 3336) | 3.73 (0.08) | |
| sheet (N = 0) | --- (---) | |
| loop (N = 480) | 0.77 (0.26) | |
| Rotamer outliers (%) | 3.68 | |
| C$\beta$ outliers (%) | NA | |
| Peptide plane (%) | | |
| Cis proline/general | 0.0/0.0 | |
| Twisted proline/general | 0.0/0.0 | |
| CaBLAM outliers (%) | 0.00 | |
| ADP (B-factors) | | |
| Iso/Aniso (#) | 32016/0 | |
| min/max/mean | | |
| Protein | 12.13/73.95/32.59 | |
| Nucleotide | --- | |
| Ligand | --- | |
| Water | --- | |
| Occupancy | | |
| Mean | 1.00 | |
| occ = 1 (%) | 100.00 | |
| 0 < occ < 1 (%) | 0.00 | |
| occ > 1 (%) | 0.00 | |
| **Data** | | |
| Box | | |
| Lengths (Å) | 130.67, 130.67, 130.67 | |
| Angles (°) | 90.00, 90.00, 90.00 | |
| Supplied Resolution (Å) | 2.0 | |
| Resolution Estimates (Å) | Masked | Unmasked |
| d FSC (half maps; 0.143) | --- | --- |
| d 99 (full/half1/half2) | 2.2/---/--- | 2.1/---/--- |
| d model | 2.2 | 2.2 |
| d FSC model (0/0.143/0.5) | 2.0/2.0/18.2 | 2.0/2.1/21.4 |
| Map min/max/mean | -0.17/0.30/0.00 | |
| **Model vs. Data** | | |
| CC (mask) | 0.82 | |
| CC (box) | 0.67 | |
| CC (peaks) | 0.65 | |
| CC (volume) | 0.80 | |
| Mean CC for ligands | --- | |

**Table 7: Deposited Structures:** Here all deposited structures are listed with their corresponding PDB-IDs.

| Name | PDB-ID |
|---|---|
| VgrG1-Spy | 28YN |
| $\beta$-galactosidase-ALFA | 28YO |
| Spy-ApoF | 28YP |
| ALFA-ApoF | 28YQ |
| ALFA-ApoF from IVT | 28YR |

#### 1.7 SI-7: Isolation of ALFA-ApoF from IVT

During our experiments, we observed that multimeric proteins like ApoF need time to assemble, otherwise mostly monomers or incomplete assemblies are isolated (see Figure 27).

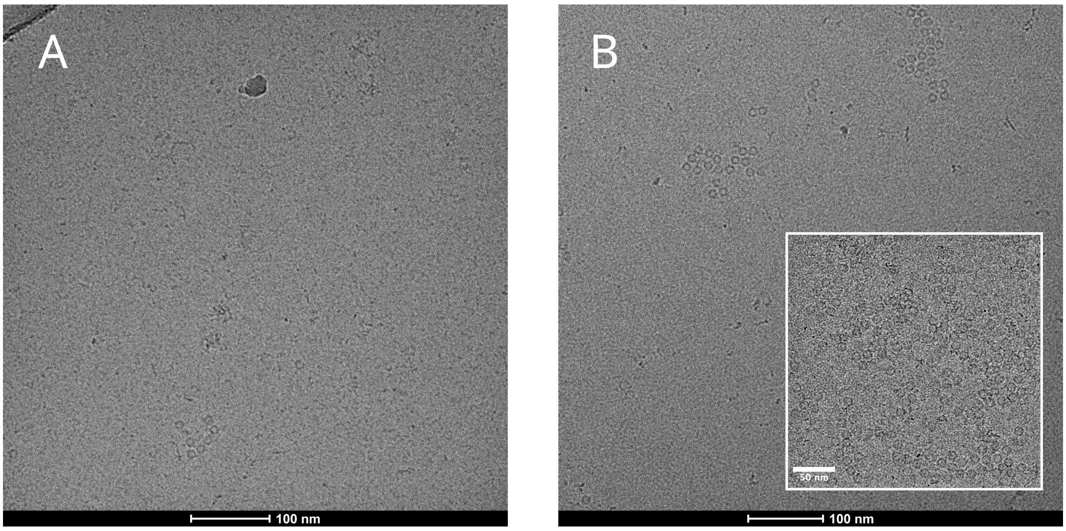

**Figure 27: Isolation of ALFA-ApoF from IVT-sample: a)** Isolation from IVT-sample immediately after IVT-reaction. Some ApoF particles are present, but a lot of smaller particles (4-5 nm) are much more abundant. **b)** Isolation from IVT-reaction where the sample was incubated at 4 °C overnight. More ApoF particles are visible (see inset image) and basically no small particles were present in the ice (see large image) suggesting that ApoF needs time to assemble into its oligomeric form. Images taken on Talos 200 kV; inset image taken during data collection on Titan G4 300 kV.

The expression of ALFA-ApoF in the IVT-reaction was confirmed by SDS-PAGE, prior to the microfluidic isolation experiments. A faint additional band around 20 kDa can be seen compared to the control reaction (see Figure 28).

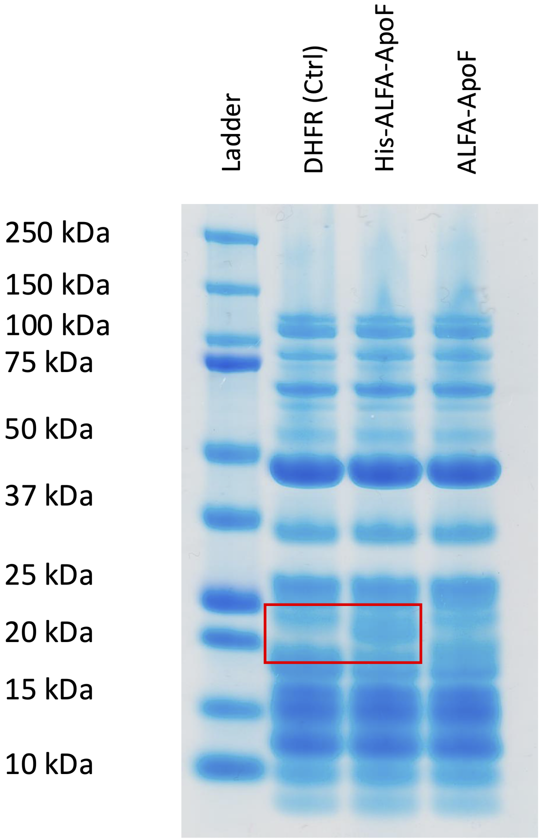

**Figure 28: SDS-PAGE gel of ALFA-ApoF IVT-reaction:** ALFA-ApoF was produced *in vitro*. An additional band around 20 kDa, which corresponds to ALFA-ApoF, can be seen compared to the control reaction where Dihydrofolate reductase (DHFR) was produced.
